# Arabidopsis leaf hydraulic conductance is regulated by xylem-sap pH, controlled, in turn, by a P-type H^+^-ATPase of vascular bundle sheath cells

**DOI:** 10.1101/234286

**Authors:** Yael Grunwald, Noa Wigoda, Nir Sade, Adi Yaaran, Tanmayee Torne, Sanbon Chaka Gosa, Nava Moran, Menachem Moshelion

## Abstract

The leaf vascular bundle sheath cells (BSCs) that tightly envelop the leaf veins, are a selective and dynamic barrier to xylem-sap water and solutes radially entering the mesophyll cells. Under normal conditions, xylem-sap pH of <6 is presumably important for driving and regulating the transmembranal solute transport. Having discovered recently a differentially high expression of a BSCs proton pump, AHA2, we now test the hypothesis that it regulates this pH and leaf radial water fluxes.
We monitored the xylem-sap pH in the veins of detached leaves of WT Arabidopsis, AHA mutants, and *aha2* mutants complemented with *AHA2* gene solely in BSCs. We tested an AHA inhibitor and stimulator, and different pH buffers. We monitored their impact on the xylem-sap pH and the whole leaf hydraulic conductance (K_leaf_), and the effect of pH on the water osmotic permeability (P_f_) of isolated BSCs protoplasts.
Our results demonstrated that AHA2 is necessary for xylem-sap acidification, and in turn, for elevating K_leaf_. Conversely, knocking out AHA2 alkalinized the xylem-sap. Also, elevating xylem sap pH to 7.5 reduced K_leaf_ and elevating external pH to 7.5 decreased the BSCs P_f_.
All these demonstrate a causative link between AHA2 activity in BSCs and leaf radial water conductance.

**One-sentence summary:** Bundle-sheath cells can control the leaf hydraulic conductance by proton-pump-regulated xylem sap pH

## Introduction

The majority (95%) of nutrients and water which enter the plant through the roots move upward in a bulk flow (“transpiration stream”) via xylem vessels all the way into the leaf veins (Taiz and Zeiger, 2014), then radially, through the bundle sheath cells (BSCs) – into the leaf photosynthetic tissue, the mesophyll. The parenchymatous BSCs, tightly enwraping the entire vasculature (Kinsman and Pyke, 1998), constitute a selective barrier between the vein and the mesophyll. For example, they impeded the transport of sodium (Shapira et al., 2009) and of boron (Shapira et al., 2013) in a banana leaf, and boron – also in Thelungiella (Lamdan et al., 2012) and Arabidopsis (Shatil-Cohen and Moshelion, 2012). Importantly, the BSCs act also as a barrier to the passage of water (Shatil-Cohen et al., 2011; Pantin et al., 2013). However, it is still not clear (Geilfus, 2017) whether and how the apoplastic (xylem) milieu modulates the “barrier behavior” of the bundle sheath, and in particular, whether this behavior depends on the xylem sap pH.

Some direct determinations of leaf xylem pH placed it at 6.3-6.7 in cotton leaves (Hartung et al., 1988), or at 5.3 in sunflower leaves (Jia and Davies, 2007) but most frequently, under normal growth conditions the leaf apoplast pH is around 5.5-6 (reviewed by Grignon and Sentenac, 1991; Geilfus, 2017). The leaf xylem sap pH can change in response to changes in external conditions experienced by the root. For example, the tomato leaf xylem sap alkalinized upon nitrate supply to roots (Jia and Davies, 2007). Barley leaf sub-stomatal apoplast also alkalinized upon roots exposure to salts (KCl, NaCl, NaNO3 (Felle et al., 2005)). Drying soil in tomato (Jia and Davies, 2007) and in *Vicia faba* (Karuppanapandian et al., 2017) or cold stress at the root level in barley (Felle et al., 2005) also caused alkalinization (measured, respectively, in the tomato xylem sap, in the *V. faba* xylem sap and leaf apoplast, and in the barley sub-stomatal apoplast). Finally, alkalinization of the barley leaf sub-stomatal apoplast was brought about also by applying ABA (considered not only a drought-stress hormone, but also shown to mediate cold stress (Huang et al., 2017)) both to barley roots and to detached leaves (Felle et al., 2005).

In turn, changes in the leaf xylem sap pH regulate physiological processes. Feeding high-pH solutions to detached leaves reduced stomatal conductance and transpiration both in *Commelina communis* and in tomato (Wilkinson and Davies, 1997; Wilkinson et al., 1998; Jia and Davies, 2007). As pH defines the dissociation state of weak acids, among them the major phytohormones abscisic acid (ABA) and indol acetic acid (IAA, auxin), changes in pH would affect both their distribution between the apoplast and the cellular compartments of the plant, and hence, their biological activities. For example, ABA accumulated in the alkalinizing xylem sap of a gradually pressure-dehydrated detached cotton leaf, which could be prevented by pretreatment with an H^+^-ATPase-activating fungal toxin, fusicoccin (Hartung et al., 1988). Leaf xylem alkalinization in *Vicia faba* as a result of gradually drying soil was followed by elevated ABA levels in the xylem (even if with a few days delay (Karuppanapandian et al., 2017).

Already widely-accepted is the crucial role of apoplastic protons in the proton-motive force governing transmembrane transport (Serrano, 1988; Haruta and Sussman, 2012; Taiz and Zeiger, 2014), and, therefore, by extrapolation, the role of xylem sap pH – in *driving* the transport between the xylem and the surrounding BSCs (Shapira et al., 2009). Xylem pH may have also *regulatory* effects on the transport proteins in the BSCs membrane, including aquaporins. Surprisingly, however, even though aquaporins largely determine the osmotic membrane water permeability coefficient, P_f_ (see also Discussion), and while P_f_, particularly the P_f_ of the BSCs, may well be important in determining the hydraulic conductance (K_leaf_) of the entire leaf (Shatil-Cohen et al., 2011; Sade et al., 2014; suggested also by modeling, wherein KOX, the lumped Outside-Xylem hydraulic conductance, is equivalent to our definition of K_leaf_ (Scoffoni et al., 2018)), the effect of *extracellular* pH (or, rather, the lack thereof) has been mentioned only in passing in relation to the P_f_ of plasma membrane vesicles of *Beta vulgaris* storage root (Alleva et al., 2006) and was ignored in relation to tobacco aquaporins (NtPIP2;1 and NtPIP1) expressed in yeast (Fischer and Kaldenhoff, 2008). Thus, the effect of apoplastic pH on P_f_ in other plant cells, and especially the effect of xylem sap pH on the BSCs P_f_, is unknown. Furthermore, in spite of the importance of apoplast pH regulation to the plant basic life processes (Haruta and Sussman, 2012), molecular evidence linking the leaf xylem sap regulation to a specific H^+^-ATPase and to the ensuing physiological changes in the leaf is missing.

H^+^-ATPases constitute a family of proton pumps driven by hydrolysis of ATP and are found in the plasma membrane of plants and fungi (Axelsen and Palmgren, 2001). Among the 11 H^+^-ATPases isoforms reported in Arabidopsis, the AHA1 and AHA2 are by far the most abundantly expressed members of this family throughout plant life and tissues (Haruta et al., 2010). Both AHA2 (Wang et al., 2014) and AHA1 (Yamauchi et al., 2016) take part in stomatal opening. In addition, a GUS reporter gene under the *AHA2* promoter, revealed abundant expression in Arabidopsis, especially in the vascular tissue of roots and leaves (Fuglsang et al., 2007). Our transcriptome analysis of protoplasts isolated from Arabidopsis BSCs and mesophyll cells (MCs) showed that the BSCs express the *AHA2* gene abundantly and at a threefold higher level than the MCs, while the *AHA1* gene expression, though also abundant, was not different between these two cell types (the other 9 AHA isoforms were expressed at much lower levels in both cell types, Wigoda et al., 2017, resembling the finding in the whole plant, Haruta et al., 2010).

Here we show how the BSCs act as a dynamic barrier for water flow between the xylem and mesophyll; we demonstrate a causative link between the activity of the BSCs AHA2 and the hydraulic conductance of the leaf and reveal that its underlying mechanism is the pH-controlled osmotic water permeability of the BSCs membranes.

## Materials and Methods

### Plant material

#### Plant types

We used WT (wild type) *Arabidopsis thaliana* plants ecotype Colombia, Col-0 (aka Col) and T-DNA insertion *AHA* mutants (Col): *AHA1* mutants: *aha1-6 (*SALK_ 016325), *aha1-7 (*SALK_ 065288) and *aha1-8* (SALK_ 118350), and *AHA2* mutants; *aha2-4* (SALK_ 082786) and *aha2-5* (SALK_022010) obtained from the Arabidopsis Biological Resource Center (Ohio State University). The plants’ genotypes were confirmed by PCR with allele-specific primers (supplemental Table S1). The single-gene mutants were confirmed further for the near-absence of *AHA2 (or AHA1)* RNA using real-time PCR (RT-PCR). For single-cell P_f_ experiments, we used *SCR:GFP* (Col) Arabidopsis plants expressing GFP specifically in the BSCs ER generated in our lab as in Attia et al. (2018).

#### Plant Growth Conditions

The soil grown plants (as in (Shatil-Cohen et al., 2011)) were kept in a growth chamber under short-day conditions (10-h light) and a controlled temperature of 20–22°C, with 70% humidity. Light intensity at the plant height was 150-200 μmol m^−2^ sec^−1^ (an optimal level based on (Wu et al., 2009), achieved using either fluorescent lights (Osram cool white, T5 FH 35W/840 HE, Germany) or LED light (EnerLED 24V-5630, 24W/m, 3000K (50%) + 6000K (50%)). The plants were irrigated twice a week.

### Determination of xylem sap pH in detached leaves by fluorescence imaging

To monitor the pH within the leaf minor veins, we used the fluorescence of the membrane-impermeant, ratiometric (dual-excitation) dye, fluorescein isothiocyanate conjugated to 10 kD dextran (FITC-D) perfused via petioles (at a concentration of 100 μM) into detached Arabidopsis leaves (Hoffmann and Kosegarten, 1995; Mühling et al., 1995; Lanz et al., 1997; Pitann et al., 2009b; Pitann et al., 2009a; Suppl. Figs. S1-S2; see Supplemental Materials and methods detailing our specific approach to leaf perfusion and sample preparation for fluorescence imaging).

#### Fluorescence microscopy

was *p*erformed using an inverted microscope (Olympus-IX8 integrated within the Cell-R system, http://www.olympus-global.com), via an Olympus UPlanSApo 10*X*/0.40 (∞/0.17/FN26.5) objective. Pairs of images were recorded using a 12-bit CCD camera, Orca-AG (Hamamatsu, http://www.hamamatsu.com), at two excitation wavelengths (488 and 450 nm, applied approx. 200 ms apart (Hoffmann and Kosegarten, 1995; Mühling et al., 1995)) and a single emission wavelength (520 nm), and saved in a linear 16 bit tiff format for further processing (Suppl. Materials and methods).

#### Image analysis

The images were processed with ImageJ (ver. 1.49V; http://rsbweb.nih.gov/ij/). A pixel by pixel ratio of the two background-corrected images in each pair was calculated using the ImageJ ‘Ratio-plus’ plugin (Paulo J. Magalhães, 2004; Suppl. Materials and Methods).

#### pH calibration curve

An *in-vivo* pH calibration curve was constructed using the dye dissolved in XPS^b^ buffered to predefined pH values (5, 5.5,6, 6.5, 7 and 7.5; see Solutions below) and fed into the leaves. This calibration curve (Suppl. Fig. S2d) served to convert the mean fluorescence ratio obtained from a minor vein segment to a pH value for each leaf (biological repeat). Occasionally, we verified the system stability using an *in-vitro* calibration curve derived from imaging drops of the calibration solutions placed directly on microscope slides (Suppl. Fig. S2d).

### Generation of *SCR:AHA2*-complemented plants

#### Vector construction

*AHA2* gene was cloned into pDONR™ 221 (Invitrogene) vector and the SCR promoter into pDONRP4P1r using Gateway® compatible by BP reactions, and later cloned into the pB7M24GW (Invitrogene) two fragment binary vector by LR reaction according to the manufacturer’s instructions. The binary *SCR:AHA2* vector was transformed into Agrobacterium by electroporation; transformants were selected on LB plates containing 25 μg/mL gentamycin and 50 μg/mL spectinomycin.

#### *aha2-4 and aha2-5 mutant lines transformation* with *SCR:AHA2*

was performed using the floral dip method (Clough and Bent, 1998). Transformants were selected based on their BASTA resistance, grown on plates with MS (Murashige and Skoog, Duchefa cat# M222.0050) Basal medium + 1 % sucrose and 20 μg/mL BASTA (Glufosinate Ammonium, Sigma cat # 45520). DNA insertion was verified in selected lines by PCR targeting the junction of *AHA2* and the 35S terminator with a forward primer about 1000 bp from the 3’ end of *AHA2* and reverse primer on the 35S terminator (see primer list in supplemental Table S1).

### Determination of AHA2 and AHA1 gene expression level in the whole leaf by qRT-PCR

#### RNA extraction and quantitative real-time (qRT-) PCR

Total RNA was extracted from leaves using Tri-Reagent (Molecular Research Center, cat. #: TR 118) and treated with RNase-free DNase (Thermo Scientific™, cat. #: EN0525). Complementary DNA (cDNA) was prepared using the EZ-First Strand cDNA synthesis kit (Biological Industries cat. #: 2080050) according to the manufacturer’s instructions. qRT-PCR was performed using C1000 Thermal Cycler (Bio-Rad), in the presence of EvaGreen (BIO-RAD cat.# 172-5204) and PCR primers to amplify specific regions of the genome (Haruta et al., 2010; suppl. Table S1). The results were analyzed using Bio-Rad CFX manager™ software. Dissociation curve analysis was performed at the end of each qRT-PCR reaction to validate the presence of a single reaction product and lack of primer dimerization. Expression levels of the examined genes were normalized using two normalizing genes (AT5G12240 and AT2G07734, Wigoda et al., 2017).

### Physiological characterization of the leaf (gas exchange and hydraulic conductance, K_leaf_)

#### Sample preparation

In all K_leaf_ determinations, we followed our previous protocols (Shatil-Cohen et al., 2011; Sade et al., 2014; Sade et al., 2015), with a necessary adaptation in the present work (detailed in the Suppl. Materials and methods) due to an approx. three-fold increase in the growth chamber light intensity (to 150-200 μmol m^−2^ sec^−1^ (Wu et al., 2009)), which resulted in increased stomatal conductance and transpiration. Prior to the gas-exchange measurements, the leaves were exposed to constant ambient conditions (1.3-1.5 kPa vapor pressure deficit (VPD), 200 μmol m^−2^ s^−1^illumination and temperature of 24 °C) for 15-20 minutes. During this period the transpiration, E, reached steady state (the values of E varied on average within 4-5%).

The measurements were conducted between 10:00 AM to 1:00 PM (1-4 hours after Lights On).

#### Gas-exchange assays

to determine gs (stomatal conductance) and E (transpiration rate) were performed (as in Sade et al., 2014) using a Li-Cor 6400 portable gas-exchange system (LI-COR, USA https://www.licor.com/) equipped with a standard leaf cuvette with an aperture diameter of two cm^2^, appropriate for Arabidopsis leaves, using light intensity similar to the growth chamber conditions (illumination: 200 μmol m^−2^ s^−1^; the amount of blue light set to 10% of the photosynthetically active photon flux density, temperature: approximately 22 °C, VPD: approximately 1.3 kPa, and [CO2] surrounding the leaf: 400 μmol mol^−1^). Readings were recorded at a steady-state E, 3-5 min after the leaf was clamped in the Li-Cor chamber.

#### Measuring the leaf water potential, Ψ_leaf_

Immediately following the gas exchange measurement, the leaf was transferred to a pressure chamber (ARIMAD-3000; MRC Israel) equipped with a home-made silicon adaptor especially designed to fit Arabidopsis petioles into the chamber’s O-ring. Ψleaf was determined as described earlier (Sade et al., 2014).

#### Determination of the leaf hydraulic conductance, K_leaf_ in detached leaves

K_leaf_ was calculated for each individual leaf as follows (Martre et al., 2000):

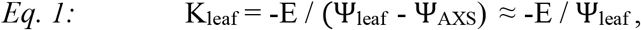

where E is the whole-leaf’s transpiration, i.e., the water flux through the leaf, Ψ_leaf_ is the leaf water potential and Ψ_AXS_ is the water potential of AXS (or of the XPS^db^); as Ψ_AXS_ is negligible in comparison to the value of leaf water potential, Ψ_leaf_, alone was used.

#### Determination of the leaf hydraulic conductance, K_leaf_, in intact leaves in a whole plant

On the evening before measurements, one leaf from each tested plant was covered by a small light-proof plastic bag. In the morning (1-4 hours after Lights On), another, illuminated leaf of the same size, positioned closely on the same plant underwent gas exchange measurements by Licor 6400 while still attached to the plant, then it was excised with a scalpel and was immediately placed in a pressure chamber to determine its water potential (as described above), taken to represent the water potential of transpiring illuminated leaves of the whole plant (Ψ_Iluminated Leaf_). The water potential of the darkplastic-covered leaf – taken to represent the water potential within the plant stem (Ψ_Dark_ Leaf =Ψ_Stem_) – was determined in an identical procedure just before the illuminated leaf. K_leaf_ was then calculated from:

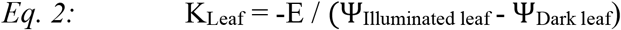

### Osmotic water permeability coefficient (P_f_) determination

#### Protoplast isolation for P_f_ experiments

Protoplasts were isolated from leaves (with verified GFP-fluresceing-veins) of 6- to 8-week-old plants with GFP-labeled BSCs (see ‘plant material’ above) by the rapid and low-stress method of (Shatil-Cohen et al., 2014) using an isotonic solution in the extraction process (pH 6, 600 mOsmol, see Solutions below). After isolation, only cells expressing GFP – i.e., only BSCs – were assayed for P_f_ determination.

#### P_f_ determination

P_f_ was determined as described by Shatil-Cohen et al. (2014), except here we used an inverted epifluorescent microscope (Nikon eclipse TS100) with a 20x/NA 0.40 objective (Nikon) and a CCD 12 bit camera Manta G-235B (https://www.alliedvision.com), and an image-processing software AcquireControl® v5.0.0 (https://www.alliedvision.com). We recorded the BSCs swelling in response to hypo-osmotic challenge (of 0.37 MPa) generated by changing the bath solutions from isotonic (600 mOsm) to a hypotonic one (450 mOsm, ibid.). The challenges were performed at pH 6 and at pH 7.5. The osmolarity of these solutions (variants of the XPS^db^; see Solutions) was adjusted with D-sorbitol and was verified within 1% of the target value using a vapour pressure osmometer (Wescor). P_f_ was determined by fitting the timecourse of the cell swelling using an offline curve-fitting procedure of the P_f_Fit program, as described in Shatil-Cohen et al., (2014) and detailed in Moshelion et al., (2004). We present here the values of the intital P_f_ (P_f_i__) obtained by fitting model 5 (ibid.).

### Solutions

#### FITC-D dye

(Sigma cat. #: FD10S) was added from a 10 mM stock in water (kept aliquoted, protected from light, at −20 °C) to all the XPS solutions (except in the leaf-tissue-autofluorescence (background) assays) to a final conc. of 100 μM.

#### XPS, basal Xylem Perfusion Solution

1 mM KCl, 0.3 mM CaCl2 and 20 mM D-sorbitol (Sigma cat# 8143). Upon preparation, the pH of this solution was 5.6 −5.8 and its osmolarity was approx. 23 mOsm/L.

#### Low-K^+^ XPS

the same as XPS above (used in experiments of Figs. 1, 2, 3b, Suppl. Figs. S1, S2a-c).

**FIGURE 1.**
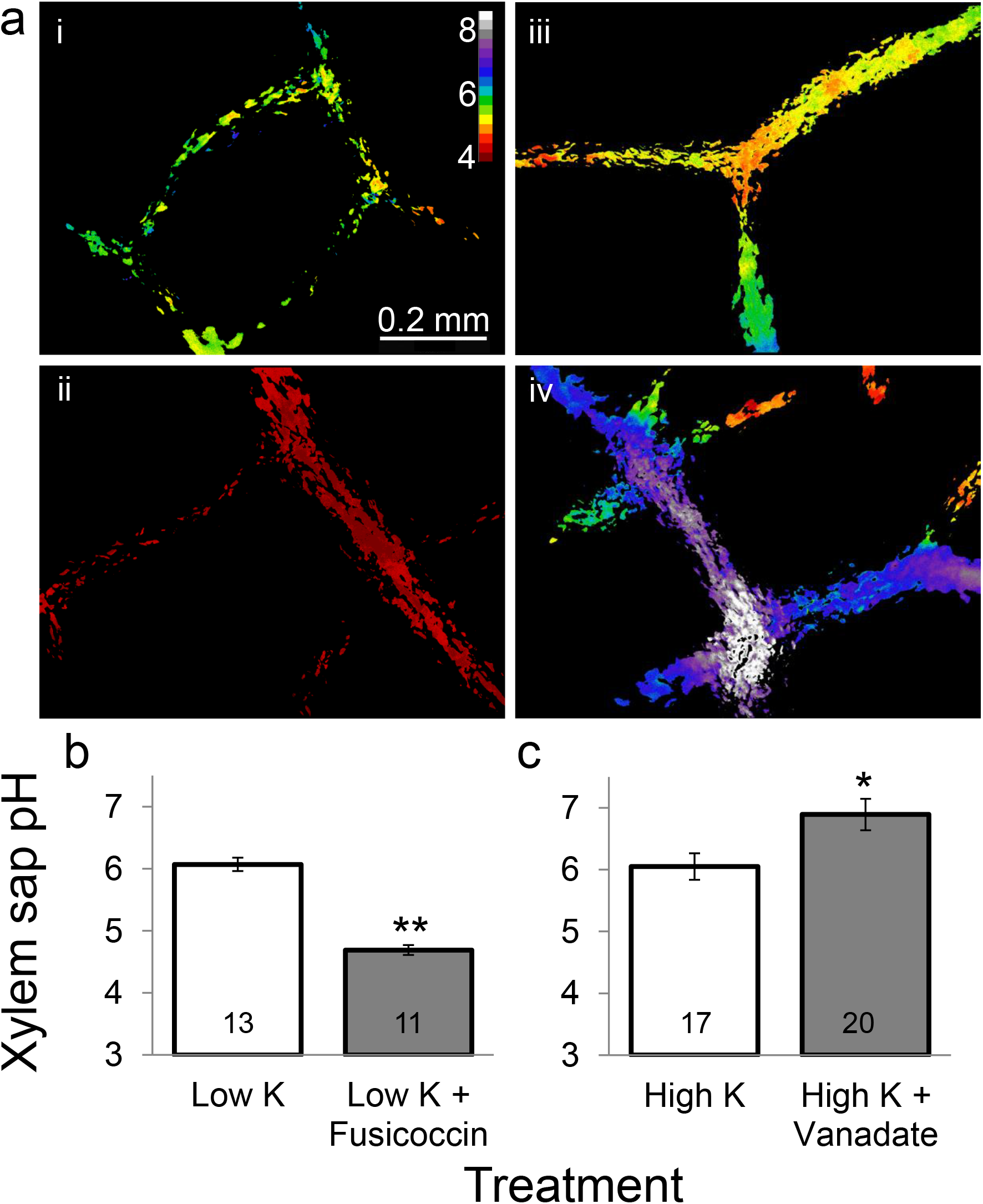
Leaf xylem sap pH reduced by petiole-fed fusicoccin (an AHA stimulator), and increased by vanadate (an AHA inhibitor) in minor veins in detached leaves of WT (wild type) Arabidopsis. **a.** Representative images of treatment effects, with color-coded pH values calculated ratiometrically for all pixels which passed the selection criteria (Materials and methods), with black masking all other pixels. i, low-K^+^ control (i.e., XPS, xylem perfusion solution). ii, XPS + 10 μM fusicoccin. iii, high-K^+^ XPS (XPS + 10 mM KNO3). iv, high-K^+^ XPS + 1 mM vanadate (Na_3_VO_4_). **b.** Mean (±SE) xylem sap pH without and with fusicoccin in the indicated number of leaves (biological repeats) from at least three independent experiments. Asterisks denote significant differences from the respective control, using Student’s two-tailed unpaired t-test (*: *P* < 0.05, **: *P* < 0.001). Other details as in a. **c.** Mean (±SE) xylem sap pH without and with vanadate. Other details as in b.

**FIGURE 2.**
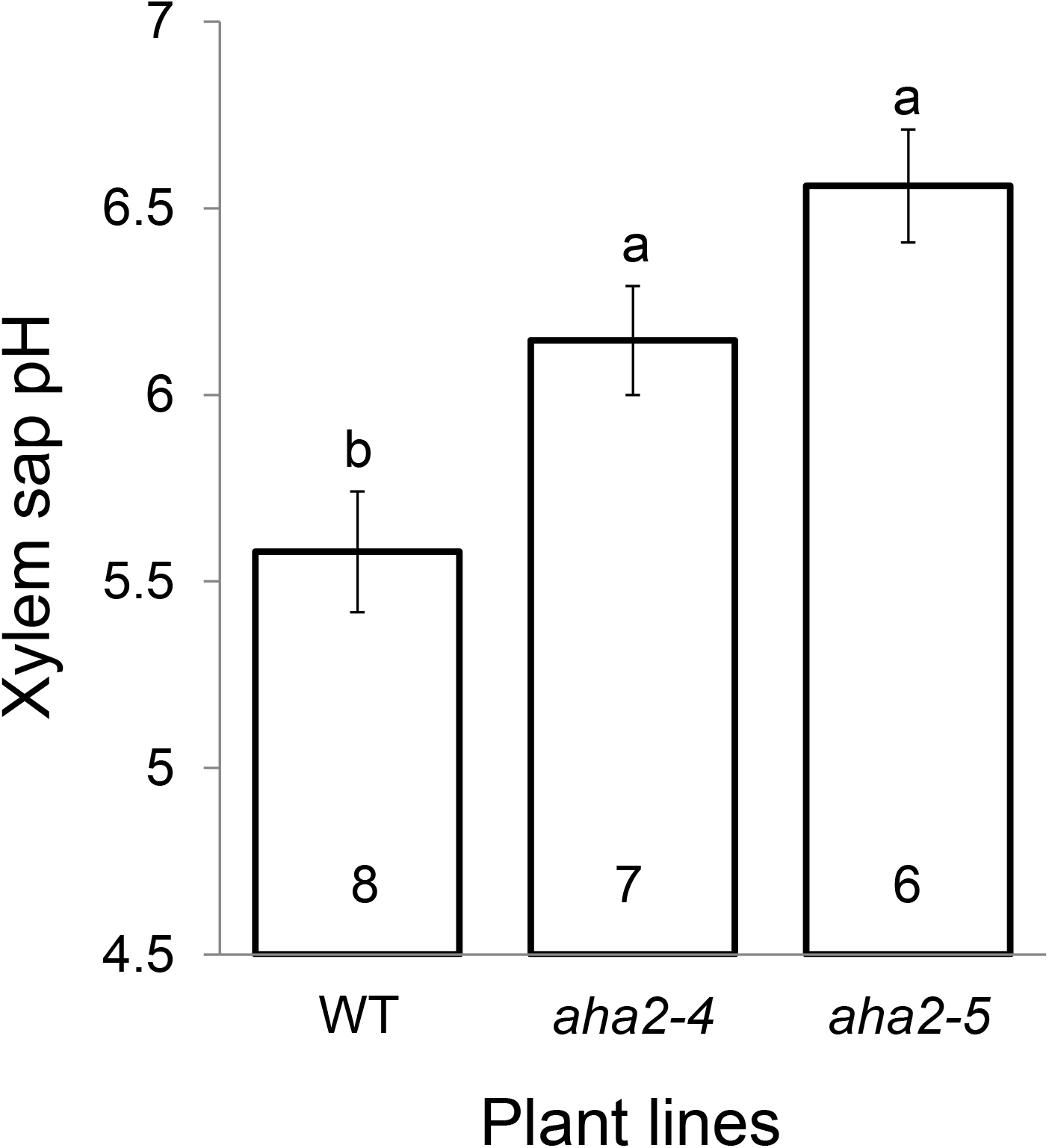
Knockout of *AHA2* increases xylem sap pH in minor leaf veins of Arabidopsis leaf. *aha2* knockout lines and WT plants. The mean (±SE) xylem sap pH in the indicated number of leaves from three independent experiments. Different letters denote significantly different pH values (P< 0.05; ANOVA).

**FIGURE 3.**
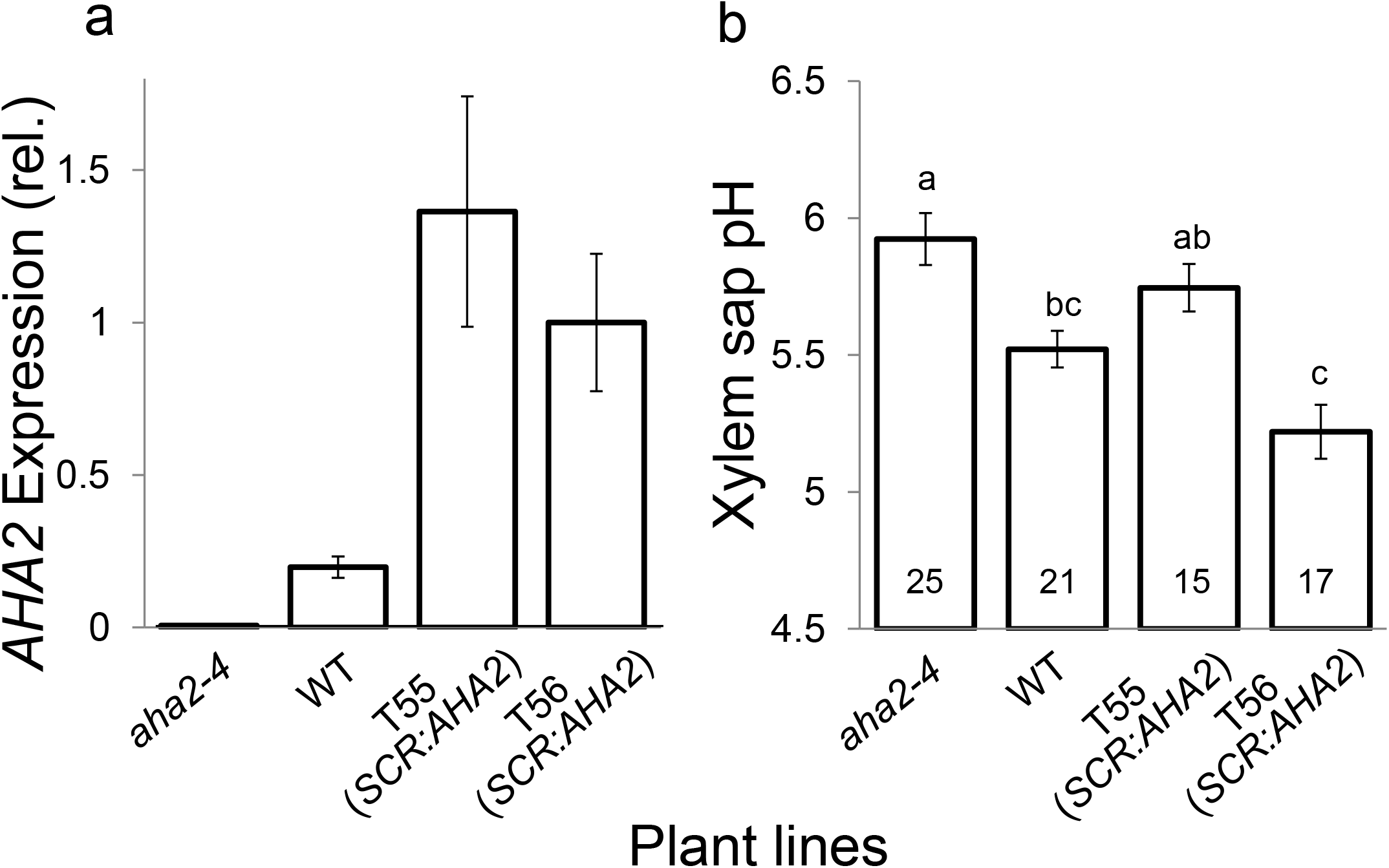
AHA2 expression lowers the xylem sap pH. Expression levels of *AHA2* and xylem sap pH in leaves of WT, *aha2-4* and of two independent lines (T55, T56) of *aha2-4* complemented with *AHA2* under a bundle-sheath-specific promotor *(SCR:AHA2)*. **a.** Mean normalized (±SE) *AHA2* expression levels obtained by qRT-PCR on whole leaf RNA (n = 5 biological repetitions, leaves); **b.** Mean (±SE) xylem sap pH in the indicated number of leaves from three independent experiments. Different letters indicate significantly different means (*P*<.0.05; ANOVA).

#### High-K^+^ XPS

XPS + 10 mM KNO3 and D-sorbitol adjustment to approx. 23 mOsm/L. Upon preparation, when unbuffered, the pH of this solution was 5.6-5.8 (used in experiments of Fig. 1).

#### XPS^b^ (pH calibration solutions)

XPS buffered with 20 mM MES and HCl or NMG (N-Methyl D-glucamine) to pH 5, 5.5, or 6), or with 20 mM HEPES and NMG to pH 6.5, 7, or 7.5. Osmolarity of these solutions was adjusted with D-sorbitol to 23 mOsm/L (used in experiments of Suppl. Fig. 2d-e).

#### AXS (*for leaf physiological characterization*)

3 mM KNO_3_, 1 mM Ca(NO_3_)_2_, 1 mM MgSO_4_, 3 mM CaCl_2_, 0.25 mM NaH_2_PO_4_, 90 μM EDFS (Sigma) and a micro-nutrient mix of 0.0025 μM CuSO_4_, 0.0025 μM H_2_MoO_4_, 0.01 μM MnSO_4_, 0.25 μM KCl, 0.125 μM H3BO3, 0.01 μM ZnSO_4_; 21 mOsm; pH of the non-buffered solution was 5.8 (used in experiments of Fig. 4a and Suppl. Figs. S3a, S3c, S3e).

**FIGURE 4.**
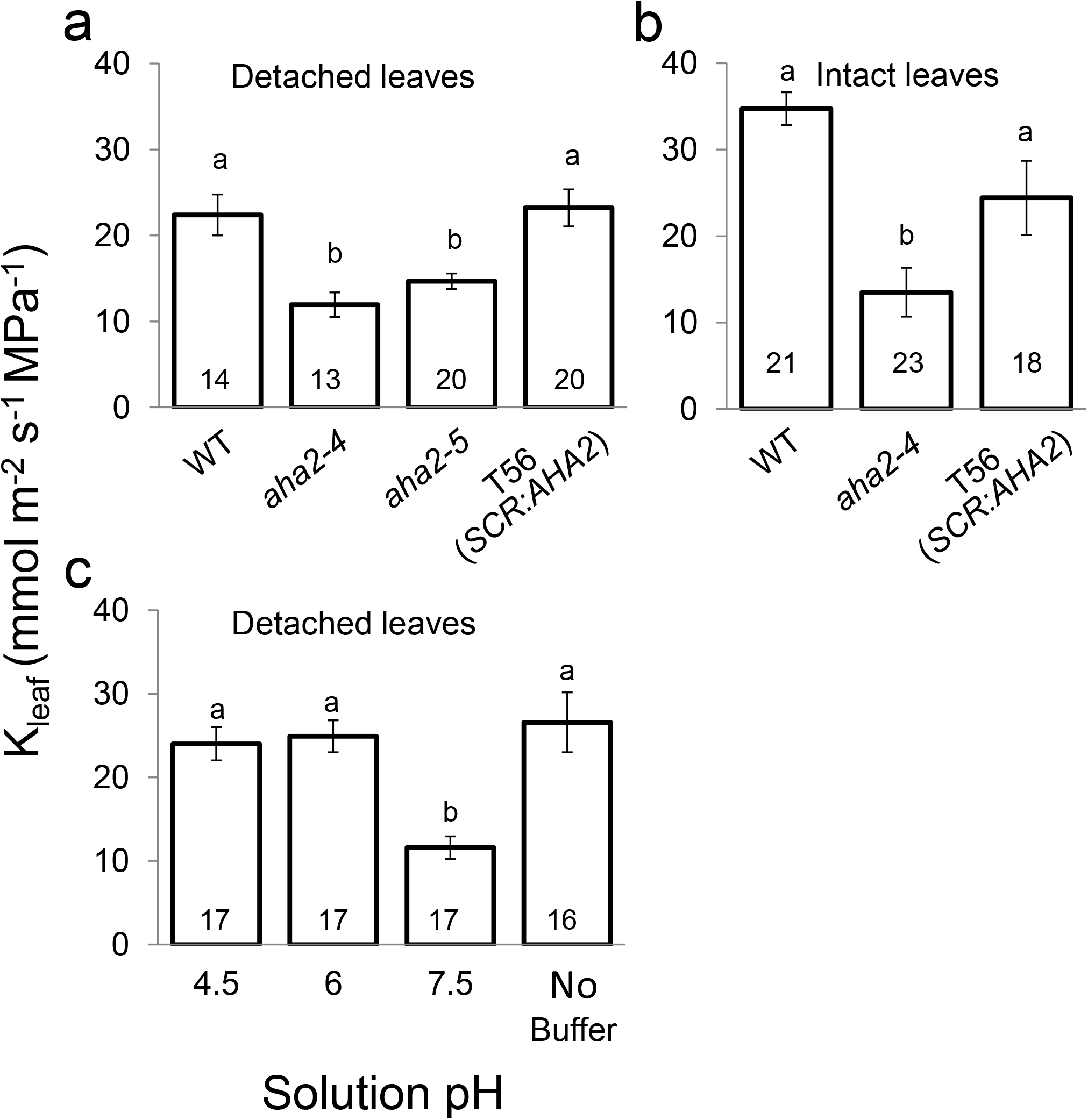
The leaf hydraulic conductance (K_leaf_) depends on the activity of AHA2 and the pH of the xylem perfusion solution (XPS). Mean (±SE) K_leaf_ (calculated by Eq. 1, Materials and Methods). **a.** K_leaf_ in detached Arabidopsis leaves of WT, *AHA2* mutants *(aha2-4, aha2-5)* and *AHA2*-complemented *aha2-4* mutant (line T56 *(SCR:AHA2))* plants perfused with non-buffered AXS. Numbers are those of assayed leaves from three independent experiments. Different letters indicate significantly different means (*P*<.0.05; ANOVA). **b.** K_leaf_ determined in intact leaves of whole Arabidopsis WT, *aha2-4* and T56 *(SCR:AHA2)* plants. Other details as in a. **c.** K_leaf_ measured in detached WT Arabidopsis leaves fed with XPS^db^ buffered to the indicated pH or with high-K^+^ non-buffered XPS. Other details as in a. Note the lower K_leaf_ at the alkaline pH.

#### XPS^db^ (*for leaf physiological characterization*)

High-K^+^ XPS buffered with 5 mM MES + 5 mM HEPES, and adjusted to pH 4.5, 6 or 7.5 with HCl or NMG. Each solution was adjusted with D-sorbitol to a final osmolarity of 32 mOsm/L (used in experiments of Fig. 4c and Suppl. Figs. S3b, S3d, S3f).

#### Solutions for P_f_ determination

pH 6 or pH 7.5 solutions: XPS^db^ adjusted to pH 6 or pH 7.5 by NMG, and adjusted with D-sorbitol to either 450 mOsmol (hypotonic) or 600 mOsmol (isotonic).

#### Solution for protoplasts isolation

pH 6 isotonic solution as for P_f_ determination.

All chemicals unless mentioned otherwise were from Sigma Israel.

### Statistics

The Student’s unpaired two-tailed t-test was used for comparison of two means, which were considered to be significantly different at P < 0.05. For comparisons of three or more population means we used ANOVA, all-pairs Tuckey HSD (JMP® Pro 13), wherein different letters represent statistical differences at P<0.05. Images which yielded extreme ratio values which were more than 2.5 SD away from the mean were discarded. Curve fitting statistics was evaluated with ANOVA (OriginPro 2020, from OriginLab Corporation, USA).

## Results

### The effect of pharmacological agents – a pump stimulator and an inhibitor – on xylem pH

To assay the activity of AHA proton pumps in the xylem-facing membranes of bundle sheath cells (BSCs) lining the Arabidopsis leaf veins, we applied fusicoccin, a known fungal stimulator of P-type proton pumps (Serrano, 1988). When 10 μM fusicoccin was fed into the xylem, we observed a sharp decrease in xylem sap pH of roughly 1.5 pH units compared with leaves in control conditions (Figs. 1a-i, 1a-ii, 1b) approximately 30 minutes after addition of fusicoccin. ETOH, the fusicoccin solvent, by itself, had no impact on xylem pH (Suppl. Fig. S4).

Further, WT leaves treated with 1 mM of vanadate (a commonly used P-type H^+^-pump inhibitor (reviewed by Palmgren, 2001), in a high-K^+^ solution (10 mM KNO3), resulted in xylem sap alkalization of about one pH unit within 30-40 min (Figs. 1a-iii, 1a-iv, 1c). Interestingly, neither a similar exposure to vanadate in the low-K^+^ solution (XPS without added KNO_3_, suppl. Fig. S5a), nor the high-K^+^ solution by itself (Suppl. Fig. S5b) had any effect on the xylem sap pH.

### AHA2 acidifies the xylem pH

In an attempt to resolve between the relative contributions of the two abundant H^+^-ATPAses of the BSCs, AHA1 and AHA2 (Wigoda et al., 2017), to the acidification of the xylem pH, we compared the xylem sap pH of WT plants to that in T-DNA-insertion mutants of either pump. We used two independent mutant lines with a homozygous loss of function of the *AHA2* gene *(aha2-4* and *aha2-5)* and three such lines with mutated *AHA1 (aha1-6, aha1-7, aha1-8*). The xylem sap pH of both *AHA2* mutants, *aha2-4* and *aha2-5,* was consistently higher, by 0.5-1 pH units, compared to the WT plants (Fig. 2). We did not find a significant difference between the xylem sap pH in WT vs. the three lines of AHA1 mutants (Suppl. Fig. S6).

To further test the ability of AHA2 to acidify the xylem sap pH, we complemented the *AHA2* deficient plants (the *aha2-4* mutant), with the *AHA2* gene directed specifically to the BSCs (under the BSCs-specific promotor Scarecrow, SCR; Wysocka-Diller et al., 2000). Using qRT-PCR on whole leaves of mature transgenic plants, we confirmed *AHA2* absence in the *aha2-4* mutant (Fig. 3a, as in Haruta et al., 2010), and demonstrated successful complementation in two lines (T55 and T56). The transcript level of *AHA2* in these lines was even higher than in the WT (Fig. 3a). While the xylem sap pH in the *AHA2-* mutant, *aha2-4*, was significantly more alkaline than in WT plants (repeating the results of Fig. 2a above), upon the complementation of the *aha2-4* mutant with the BSC-directed *AHA2,* the xylem sap pH in both lines became no higher than WT (Fig 3b). The level of *AHA1* transcript in the whole leaf was not affected by the genetic manipulations of AHA2, neither by *AHA2* mutation (as in Haruta et al., 2010), nor by the BSC-specific complementation of the mutant with *AHA2* (Supl. Fig. S7).

### Elevating pH lowers the leaf hydraulic conductance, K_leaf_

To examine the impact of AHA2 on the water economy of the whole leaf we determined the leaf water potential (Ψ_leaf_) and its transpiration (E) at steady-state, and from these two we calculated the leaf hydraulic conductance (K_leaf_, Eq. 1 in Materials and methods).). We compared the K_leaf_ in detached leaves of WT, of the *aha2-4* and *aha2-5* mutants (respectively, 22.4, 12.0 and 14.7 mmol m^−2^ s^−1^ MPa^−1^), and of *aha2-4* complemented with the BSCs-directed *AHA2* (i.e., *AHA2* under the Scarecrow promoter; line T56, which presented the lowest xylem pH of all genetic lines), all fed with the non-buffered AXS. Notably, while all four leaf types appeared to transpire at a similar rate, E (Suppl. Fig. S3e; but see Discussion), the K_leaf_ of the mutants was appreciably lower than that of WT leaves, about 50% of WT in *aha2-4* and about 35 % in *aha2-5,* while K_leaf_ of the AHA2-complemented T56 line (23.2 mmol m^−2^ s^−1^ MPa^−1^) was practically as high as that of the WT leaves (Fig. 4a).

In the intact leaves in whole plants, while E also did not appear to differ among the three plant lines (Suppl. Fig. S8), K_leaf_ values were similar to those of K_leaf_ in detached leaves and we observed the same pattern: K_leaf_ of intact *aha2-4* plants was by about 60% lower than in WT (respectively, 13.5 vs. 34.7 mmol m^−2^ s^−1^ MPa^−1^), while K_leaf_ of T56 plants (24.4 mmol m^−2^ s^−1^ MPa^−1^) was not significantly different from WT (Fig 4b).

Furthermore, and again in contrast to E, which appeared unaffected (Suppl. Fig. S3f), pH treatments did alter the K_leaf_ of detached WT leaves. Perfusion with XPS^db^ buffered at pH 7.5 lowered K_leaf_ by roughly 50% compared to K_leaf_ in WT leaves perfused with XPS^db^ buffered at pH 6 or WT leaves perfused with the non-buffered XPS (respectively, 11.6, 24.9 and 26.6 mmol m^−2^ s^−1^ MPa^−1^; Fig. 4c). In contrast to K_leaf_ in the more alkaline conditions, K_leaf_ of leaves perfused with the more acidic pH 4.5 (24.0 mmol m^−2^ s^−1^ MPa^−1^) was no different than at the “normal” pH 6 (Fig. 4c).

### Alkaline pH lowers the osmotic water permeability coefficient of BSCs protoplast membrane, P_f_

In order to test the hypothesis that the pH-dependent reduction in K_leaf_ is due to a reduction in the water permeability of the BSCs membranes, we determined the osmotic water permeability coefficient, P_f_, of BSC under two pH treatments, using GFP labeled BSCs, from *SCR:GFP* plants. We verified that these plants did not differ in their physiological parameters (K_leaf_ and the underlying E and Ψ_leaf_) from the unlabeled WT plants (Suppl. Fig. S9). The mean initial P_f_ (see Materials and methods) of BSCs assayed in pH 6 was ~8.5 fold higher than that of BSCs assayed in pH 7.5. (5.39 ± 1.0 μm sec ^−1^ and 0.63 ± 0.2 μm sec ^−1^, respectively; Fig. 5).

**FIGURE 5.**
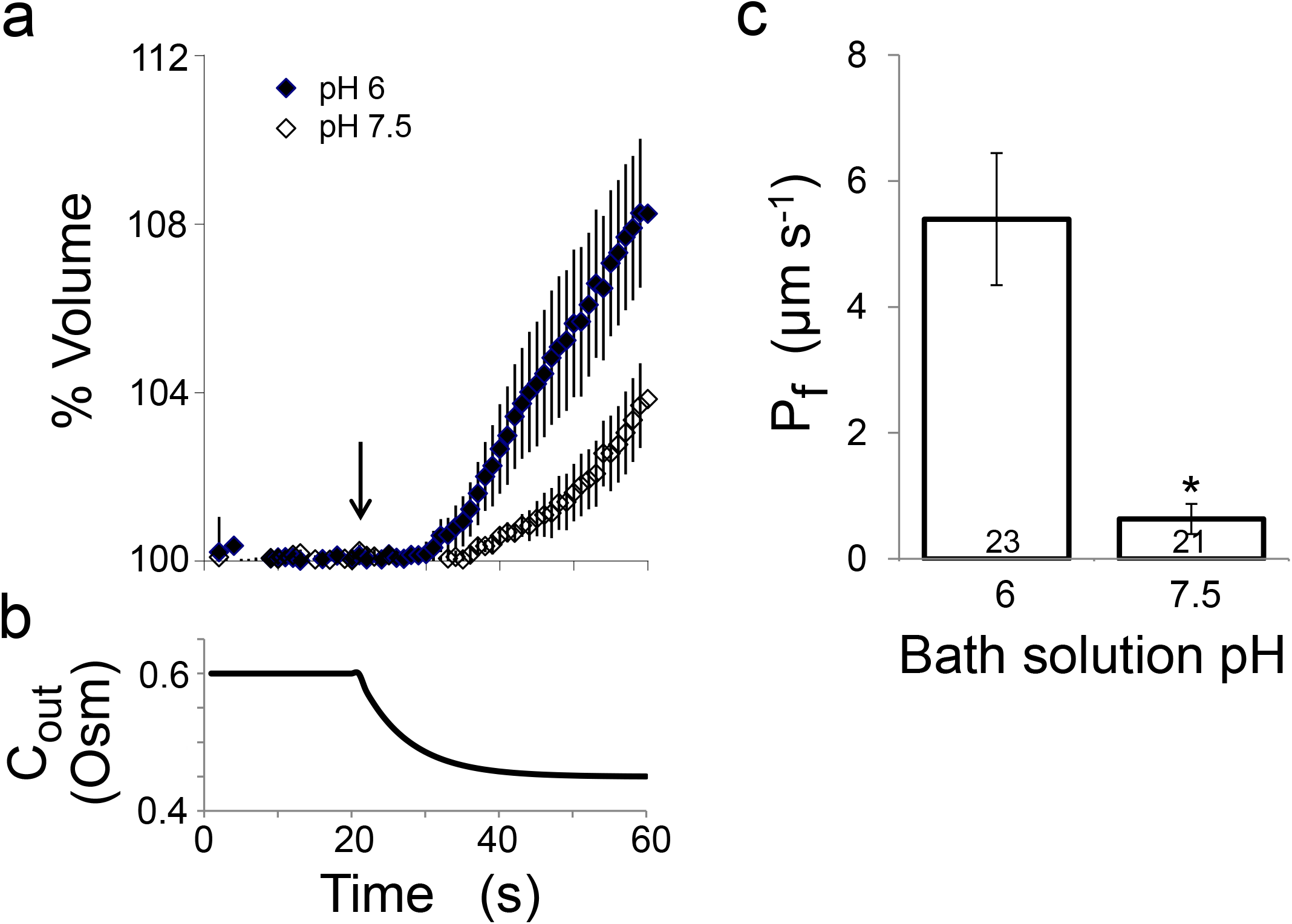
The BSCs membrane osmotic water permeability coefficient (P_f_) depends inversely on external pH. **a.** Time course (60 sec) of bundle sheath protoplasts from *SCR:GFP* plants swelling upon exposure to a hypotonic XPS^db^ solution at pH 6 or 7.5. The arrow indicates onset of bath flush. **b.** Time course of the osmotic concentration change in the bath (Cout) during the hypotonic challenge (calculated as in Moshelion et al., 2004). **c.** Mean (±SE) initial P_f_ values of the indicated number of bundle sheath protoplasts at the different pHs from three independent experiments. The asterisk denotes a significant difference between the treatments using Student’s two-tailed unpaired t-test (P < 0.01).

### Leaf veins architecture

We tested the hypothesis that K_leaf_ values are proportional to the area of the bundle sheath enwrapping the leaf veins, and therefore to the combined length of leaf veins per unit leaf area (vein density). Thus, leaves of the *aha2-4* plants would be expected to have lower vein density than leaves of WT and *SCR:AHA2* (T56) plants. However, a comparison (Suppl. Materials and methods) yielded no difference among these lines (Fig. 6, Suppl. Fig. S10), eliminating this structural reason for differences in K_leaf_.

**FIGURE 6.**
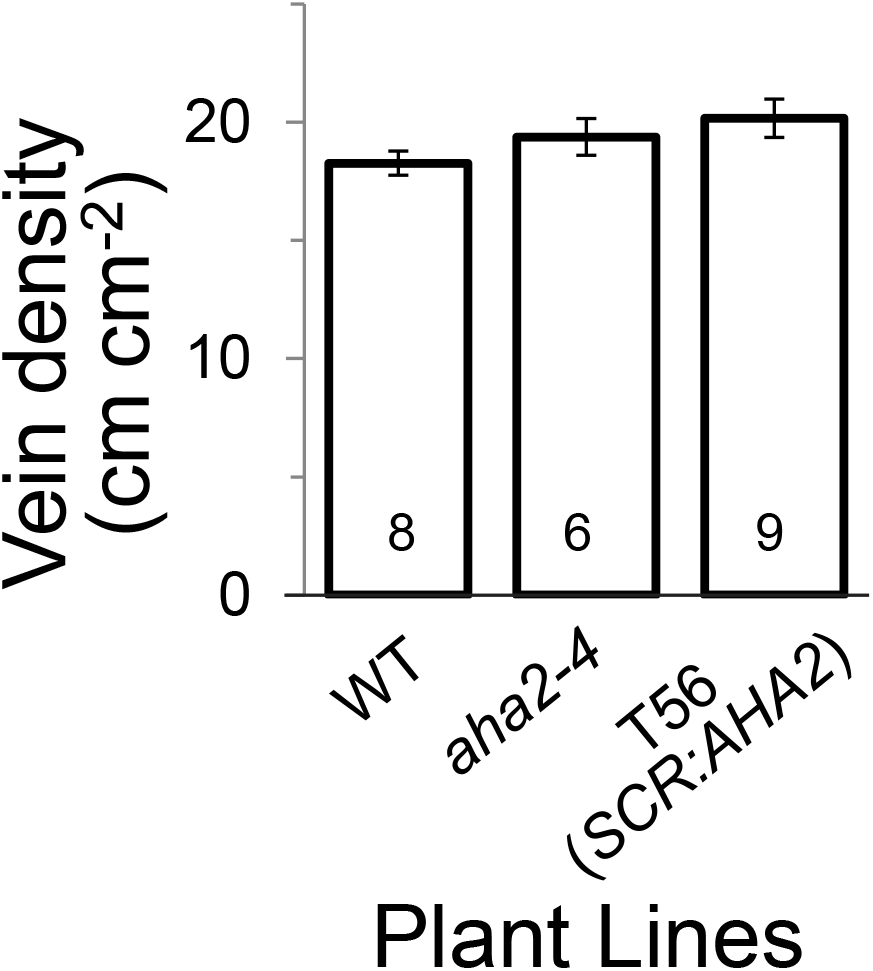
Leaf vein density does not depend on AHA2. Mean vein density (±SE) averaged over the indicated number of leaves (see details in Suppl. Fig. 10). Vein density was calculated as the total vein length divided by the scanned leaf area. Note the lack of significant difference between the three genetic lines.

## Discussion

Our conclusions from this study are based on two complementing AHA manipulation approaches: (1) classical pharmacology (using fusiccocin and vanadate) and (2), AHA2 genetics (mutants and complementation), combined with in-vivo measurements of the resulting xylem sap pH, using the petiole-fed membrane-impermeable FITC-dextran. An additional approach was to assay the physiological functions in the single-cell (P_f_), the detached leaf (K_leaf_), and the whole plant (as reflected in the *intact* leaf K_leaf_), combined with the modulation of the xylem sap pH (by genetic AHA2 manipulations or by direct pH buffering of the BSCs-facing solutions).

### The AHA2 of the BSCs is indispensable for the leaf xylem sap acidification

Plasma membrane H^+^-ATPases (AHAs) are well-known regulators of the apoplastic pH and have a key role in cell ion homeostasis. Here we demonstrate that the BSCs AHA2 exerts a dominant role in the acidification of the xylem sap in the Arabidopsis leaf.

An AHA-specific stimulator (fusicoccin) and an inhibitor (vanadate), fed directly to the leaf xylem, established that pH acidification of the leaf xylem sap involves P-type H^+^-ATPases, which extrude protons – most likely, from the BSCs – into the xylem lumen (Fig. 1). Notably, to demonstrate vanadate effect (xylem sap alkalinization) the high-K^+^ XPS was necessary, possibly to dissipate the xylem sap protons by enhanced H^+^/K^+^ (accompanied by 2H^+^/NO_3_^−^) co-transport into the BSCs. That proton-coupled K^+^ transmembrane co-transport plays a particularly important role in the BSCs is suggested by the 10% higher expression in BSCs (relative to mesophyll cells) of two K^+^ uptake permeases, AtKT2 and AtKUP11 (likely to be H^+^-coupled (Wigoda et al., 2017) and references therein). This conforms with the aforementioned general notion that the pH of an apoplastic compartment reflects, among others, a balance of bi-directional proton movements (Serrano, 1988).

While pH in the *aha1* mutant lines was not significantly different from that in WT (Suppl. Fig. S6), the pH of the xylem sap in both AHA2 mutant lines, *aha2-4* and *aha2-5* was consistently above that in WT (Fig. 2), similar to the pH elevated by vanadate inhibition (Fig. 1C). Two lines of *aha2-4* complemented with the *AHA2* gene directed specifically into the BSCs, using the specific promotor SCR, had their xylem sap pH restored to WT-like levels. These results together underlined the importance of AHA2 in determining the xylem sap pH.

### Extracellular pH regulation of the membrane osmotic water permeability

Earlier work established that low *cytosolic* pH reduces the membrane osmotic water permeability coefficient, P_f_, by inhibiting aquaporin gating (Gerbeau et al., 2002; Tournaire-Roux et al., 2003; Alleva et al., 2006; Fischer and Kaldenhoff, 2008; Leitão et al., 2012; Frick et al., 2013; Yaneff et al., 2016). This phenomenon is especially relevant to roots under stress conditions such as flooding or drought, where a change in the water hydraulic permeability of the roots is considered to be part of the plant stress defense mechanism.

External pH seemed not to affect the membrane hydraulic permeability of intact cells (Lp) of Nitella, Chara and maize, except a slight inhibition at pH around 4.5 in Nitella and in one variety of maize (Tyerman et al., 2002 and references therein). However, P_f_ sensitivity to pH has been also found in several mammalian cell types, be it cytosolic pH (Kaptan et al., 2015; Gotfryd et al., 2018), or extracellular pH (Mosca et al., 2018). Here we demonstrate, for the first time in an intact plant protoplast, the BSC, a reduction in P_f_ by *external* (xylem sap) alkalinization. This pH sensitivity differs from the *lack* of external pH effect on the right-side-out plasma membrane vesicles of *Beta vulgaris* root (Alleva et al., 2006). However, this difference is not surprising, in view of the differences between root and shoot, for example, in their opposite responses to drought: acidification in the root xylem sap and alkalinization in the shoot xylem sap (Karuppanapandian et al., 2017).

“Macroscopic” membrane P_f_ changes and, moreover, whole-organ K_leaf_ changes have been tied to changes in aquaporin activity. For example, Shatil-Cohen et al., (2011), reduced both P_f_ and K_leaf_ by a common aquaporin blocker HgCl_2_. Also, Prado et al. demonstrated that three aquaporins expressed in veins, and in particular, PIP2;1, contributed to darkness-enhanced hydraulic water conductance of the Arabidopsis leaves rosette (Prado et al., 2013). Additionally, BSCs-directed knockdown of aquaporins (using artificial microRNAs under the SCR promoter) decreased the K_leaf_ of a detached Arabidopsis leaf and, separately, the BSCs P_f_ (Sade et al., 2014). Therefore, it is interesting to compare the external pH effect we observed on the P_f_ of BSCs to the effects of external pH on aquaporins. For example, an external acidification (from pH 7.4 to pH 5), perceived via external tyrosine and histidine, *decreased* the permeability to water and glycerol of the human aquaglyceroporin, AQP7, with a half-inhibition at about pH 6 (Mósca et al., 2018); in contrast, the fungal aquaporin RdAQP1 activity (when expressed in Arabidopsis protoplasts) was enhanced at acidic external pH and inhibited at alkaline external pH (Turgeman et al., 2016).

Like RdAQP1, at least several Arabidopsis aquaporins have histidines (pka = 6.8) and/or cysteins (pKa=8.5) in their apoplast-facing loops, and also plenty of tyrosines (pKa=10). For example (even excluding tyrosines, because of their high pKa), based on uniprot membrane topology (www.uniprot.org/), PIP2;1 has exterior-facing Cys75 and His248; PIP2;2, has exterior-facing Cys73; His246; PIP2;3: Cys73, His148,-246, etc. Future work will clarify whether the P_f_ reduction in BSCs and the reduction of K_leaf_ in detached Arabidopsis leaves (see also further discussion of K_leaf_ and P_f_ relationship below) is also mediated by aquaporins and their external histidines/cysteins.

### AHA2 role in the regulation of the whole leaf water balance

The notion that AHA pumps, in general, power secondary H^+^-cotransport across cell membranes facing the apoplast is decades old (Spanswick, 1981; Serrano, 1988; Sze et al., 1999; Palmgren, 2001). The plant plasma membrane H^+^-ATPase has since been recognized as essential for plant growth (reviewed by Falhof et al., 2016). The practically sole “celebrated product” of its action to date is the proton motive force (PMF) – a gradient of protons concentration without- or in combination with an electrical gradient – which drives the movement of solutes across cellular membranes. In only a few cases has a *specific* physiological role been assigned to the plasma membrane H^+^-ATPase – mainly in stomatal physiology and in roots (reviewed by Falhof et al., 2016).

Here we show another novel aspect of transport activity *regulated* (rather than *driven)* by AHA2 – that of water fluxes from the xylem into the leaf and across the bundle sheath layer – evident as the leaf hydraulic conductance, K_leaf_. Our summary figure (Fig. 7) demonstrates this regulation by juxtaposing the separate measurements of of K_leaf_ (Fig. 4a) and xylem sap pH (Figs. 2, 3b) performed in independent experiments on detached leaves of WT plants, aha2 mutants and the *SCR:AHA2*-complemented aha2-4. This combination demonstrates the inverse dependence of K_leaf_ on the xylem sap pH, in particular roughly between pH 5.5 and 6.5 (Fig. 7). The inverse K_leaf_ - pH relationship is also demonstrable when pH is imposed directly by buffers in the xylem perfusate in the range of pH 6 to 7.5 (Fig. 4c, Fig. 7). Yet, while the absolute values of the maximum and the minimum K_leaf_ were similar in the two experiment types, there was a two-fold difference in the K_leaf_ values at the pH transition range (around pH 6): K_leaf_ was high (approximately 25 mmol m^−2^ s^−1^ MPa^−1^) in the WT leaf with the xylem sap buffered at pH 6 and it was low (approximately 12 mmol m^−2^ s^−1^ MPa^−1^) in the *aha2-4* mutant leaf with the non-buffered xylem sap dye-reported to be at pH 6 (Fig. 7). This apparent disparity could be bridged, if, due to the ongoing activity of their AHA2, the BSCs in the leaves of the WT plants faced a nearby-layer of medium more acidic than the bulk of the pH 6-buffered xylem sap. We represented such possible inadequacy of the perfusate buffer by a hand-drawn elipse (Fig. 7). The notion that the bulk pH may differ from the pH near the membrane is supported by the report of Martinière et al. (2018), who showed that the pH reported by a pH-dye in the bulk of the apoplast (here: the xylem sap) did not fully represent the pH existing in the immediate vicinity of the membrane. Bridged by the “elipse-correction”, the results of the different pH manipulations (the genetic and the imposed buffering), together attest to a causative inverse correlation between the xylem sap pH and the resulting K_leaf_.

**FIGURE 7:**
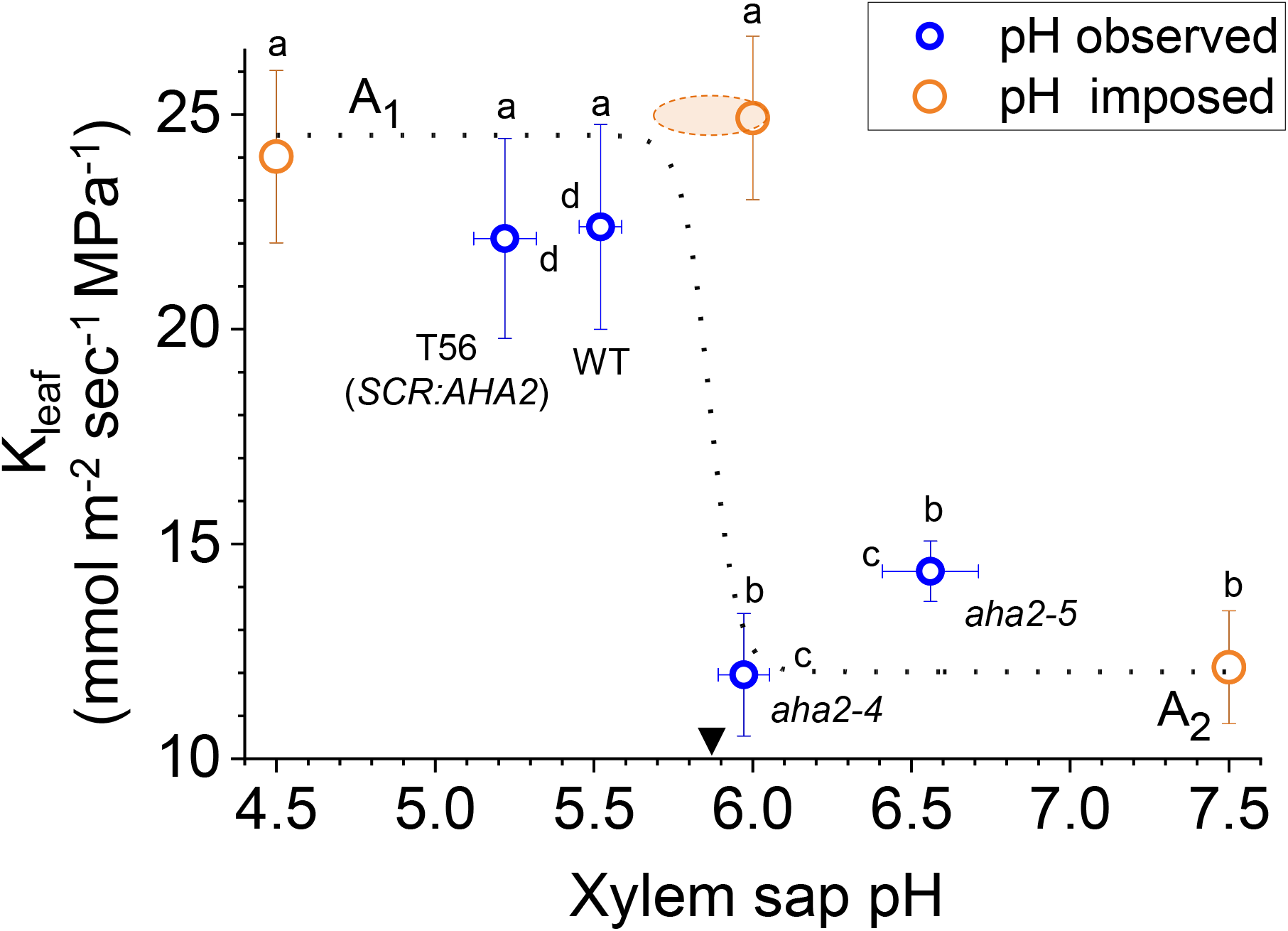
K_leaf_ depends inversely on xylem sap pH. K_leaf_ as a function of the xylem-sap pH in detached leaves. Symbols: **o,** K_leaf_ data from Fig. 4a (here: on the ordinate) vs. pH observed in detached leaves (of the same plant lines) in which the BSCs modified the pH of the unbuffered XPS or AXS within the xylem depending on AHA2 presence (and activity); the pH data are from Figs. 2 and 3b (here: on the abscissa); **o,** K_leaf_ of detached leaves of WT plants in which the xylem sap pH was imposed by perfusion with pH-buffered solutions; the data are from Fig. 4c. Different letters denote significantly different means (a, b: K_leaf_ values, c, d: pH values). Dotted line: a sigmoid K_leaf_=A_2_+(A_1_-A_2_)/(1+exp((x-x_0_)/dx)) fitted to the seven data points (orthogonal fit, OriginPro v. 2020 (9.7.0.188), Origin Lab Corporation, Northampton, USA); the maximum and minimum K_leaf_ indicated in the figure are A_1_=24.5 mmol m^2^ s^-1^, A_2_= 12.0 mmol m^−2^ s^−1^ (R^2^, the “goodness of fit” = 0.986). Note the inverse relationship between K_leaf_ and the xylem sap pH around the midpoint of x_0_ = 5.87 (arrowhead). The orange ellipse (hand drawn) describes a range of possible values of pH in the immediate vicinity of the WT BSCs, which could occur if the intact AHA2 activity in their membranes counteracted to some extent the pH of the buffered perfusate).

What is the physiological relevance of the pH-regulated K_leaf_ changes? The relatively high range of K_leaf_ values in detached leaves that we report here is characteristic of well-watered Arabidopsis plants, as reported also elsewhere (e.g., (Guyot et al., 2012; Scoffoni et al., 2018). This is exemplified in the resemblance between the relationships of the individual values of K_leaf_ vs Ψ_leaf_ (the leaf water potential) in detached and also in intact leaves (Suppl. Fig. S11) and in those of Scoffoni et al. (2018, in their Fig. 1; particularly noteworthy is the quantitative agreement between our findings and Scoffoni’s – in detached leaves – of the values of Ψ_leaf_ at which K leaf declines to 50% of its measured maximum (Scoffoni’s “P50”), −0.17 to −0.16 MPa).

Since the leaf water potential (Ψ_leaf_) reflects the relative insufficiency of water influx to the leaf in replacement for the transpired water (prior to establishing a steady-state), it is meaningful to examine the inverse relationship: how Ψ_leaf_ depends on K_leaf_. Indeed, when considering the paired measurements of Ψ_leaf_ and K_leaf_ of individual leaves, both detached and intact, a positive correlation can be demonstrated even among the well-hydrated plants (Suppl. Figs. S12a-S12c). Similarly, the rate of transpiration (E) also increases with K_leaf_ (Suppl. Figs. S12d-S12f). Incidentally, the E-K_leaf_ correlation is identical to a gS (stomata conductance)-K_leaf_ correlation, since gS=E/ VPD (Vapor Pressure Difference, the transpiration drive, which is kept constant during the Li-Cor measurements (Materials and methods)). These correlations strongly suggest that pH-controlled physiological effects – K_leaf_ and E changes – even if relatively subtle and masked by the large variability when the averages are considered, may occur in intact plants and that, in the long run, they may have cumulative effects on agricultural yield.

The above considerations of the AHA2 role invite a speculation with regard to the role of the plant stress hormone, abscisic acid (ABA) in regulating K_leaf_. In an earlier work, Shatil-Cohen et al. (Shatil-Cohen et al., 2011) localized K_leaf_ to the BSCs layer by showing that K_leaf_ decreased when the BSCs were brought into direct contact with ABA fed to the detached leaf, and not when ABA was smeared on the leaf surface. Our current work outlines a mechanism for this phenomenon: stress-induced xylem alkalinization (already reported by others (Jia and Davies, 2007; Wang et al., 2012; Korovetska et al., 2014; Karuppanapandian et al., 2017), likely due to ABA inhibition of the BSCs AHA2 (although not excluding other AHAs), similar to the H^+^-ATPase inhibition seen in guard cells (Goh et al., 1996; Schroeder et al., 2001; Zhang et al., 2004), reduces the water permeability of the BSCs membranes. Consequently, the water permeability of the entire bundle sheath layer declines, demonstrating that under stress conditions it is an active hydraulic barrier (Shatil-Cohen et al., 2011; Pantin et al., 2013). Here we show directly, using individual isolated BSCs, that P_f_, the measure of the rate of water passage via the BSCs membranes, declines with extracellular medium alkalinization (Fig. 5). Such a pH rise could mediate the effect of ABA (in a feedback loop?) on BSCs in the detached leaf.

Another, slower route of generating xylem sap alkalinization would be via decreasing the expression and/ or activity of AHA2 in the BSCs. In accord with this prediction, Genevestigator analyses (Hruz et al., 2008) of data from a few available studies, revealed that drought decreased the AHA2 expression in Arabidopsis rosettes by about 20-60% (Ludwików et al., 2009 (ArrayExpress Repository, Experiment E-MEXP-1863); Wilkins et al., 2010 (GEO repository, Expt. GSE19700)), which is in agreement with the reported alkalization of the apoplast during drought. In contrast, under drought, AHA1 expression was either unchanged (in the two above-mentioned databases) or upregulated by ~25% (Pandey et al., 2013 (GEO repository, Experiment GSE40061)), suggesting different roles for these pumps in the leaf, at least under this type of stress. Moreover, genetic manipulation of AHA2 left the quite abundant AHA1 transcript (Wigoda et al., 2017) unaffected, and AHA1 mutation left the xylem sap pH unchanged, supporting such differentiation. Notwithstanding the above findings, we have not ruled out completely a potential contribution of AHA1 to the xylem sap pH. Both AHAs need yet to be investigated in *BSCs-focused* genetic manipulations (overcoming perhaps the reported lethality of the double *aha1-aha2* mutant (Haruta et al., 2010)) and in BSCs-*specific* responses to drought.

The observed variations in transpiration rate of whole plants have been proposed to reflect internal adjustments – not only at the guard cell level but also at the K_leaf_ level – protecting the plant against challenges of changing ambient conditions (Wallach et al., 2010; Scoffoni et al., 2017; Scoffoni et al., 2018). We propose here that the dynamic adjustments in water flux through the transpiring leaf are due to the dynamic changes of BSCs’ P_f_ in response to changes of xylem sap pH and the consequent changes in K_leaf_. In a transpiring plant, K_leaf_ decreases as a consequence of xylem alkalinization in the shoot, which signals a decline of soil moisture. The K_leaf_ decrease, leading to leaf water potential decrease and consequently to stomata closure, is what prevents further water escape from the xylem and decreases the water tension in the xylem, diminishing the danger of cavitation and embolism (see also, (Guyot et al., 2012; Scoffoni et al., 2018).

How general is drought-stress-induced shoot alkalinization? While various authors demonstrated shoot xylem sap pH increase in annual plants upon gradual soil drying (for example, in sunflower from around 6 to 7 (Gollan et al., 1992); in *Commelina communis* from around 6 to over 6.5 (Wilkinson and Davies, 1997); in tomato, from 5 to 8 (Wilkinson et al., 1998); in *Vicia faba* from 5.9 to 6.8 (Karuppanapandian et al. (2017), Sharp and Davies (2009) tabulated different responses of the shoot xylem pH to drought in different perrenial plants. Therefore, in view of our work results, the leaf xylem sap pH response to water deficit and the dependence of K_leaf_ on xylem sap pH need to be examined also in other plants in order to understand the universality of the mechanism we propose.

The results of this study broaden our basic understanding of how a leaf controls its water influx supporting the notion that xylem sap alkalinization mediates the decline in plant shoot nutrient uptake due to abiotic stress. Here we focus specifically on the stress-induced cessation of activity of AHA2 in the BSCs in Arabidopsis, where we found the AHA2 instrumental in generating the low pH in the leaf xylem sap.

The consequence of this decline of AHA2 activity and xylem sap alkalinization, is a decline in the PMF. Since AHA2 – via the PMF – powers the secondary, proton-coupled export and import of solutes across the BSCs membranes, not only does it regulate plant nutrition, but, very likely also drives the extrusion of toxic compounds from the xylem-lining cells into the xylem, hence governing also the plant toxicity tolerance as well as others potential stresses. These hypotheses await future experimentation.

In conclusion, our finding that the Arabidopsis AHA2 plays a major role in regulating K_leaf_ via xylem sap pH provides a molecular basis for understanding a novel aspect, other than just PMF, of the control that the xylem sap pH can exert in the leaf. This control, in combination with the effects of PMF, is likely to underlie the plant’s key physiological activities. These results provide a new focus for exploration and understanding the role of the involvement of BSCs in determining the xylem sap composition, and, in particular, their role as a transpiration-controlling valve in series with the stomata. As the rapid growth rate and high yields of crop plants are positively correlated with enhanced transpiration and K_leaf_, BSCs are likely to become a key target tissue for the development of a new generation of manipulations for plant adaptation to environmental challenges.

## Acknowledgments

We thank Ms. Dvorah Weisman of the Hebrew University of Jerusalem for help with image analysis and Dr. Shifra Ben-Dor of the Weizmann Institute of Science for comments on the manuscript.

This research was supported by the Israel Science Foundations, ISF (grants No.1312/12, and 1842/13 to NM, grant No. 878/16 to MM and a grant No. 12-01-0007 from the Israeli Chief Scientist, Ministry of Agriculture & Rural Development to MM and NM).

## Author Contributions

### Author contributions

Y. Grunwald and N. Wigoda planned, performed and analyzed the vein pH monitoring and RT-PCR experiments, Y. Grunwald planned, performed and analyzed K_leaf_ and P_f_ experiments.Y. Grunwald and N. Wigoda wrote the paper. A. Yaaran and T. Torne participated in generating the complemented plant lines, and A. Yaaran and S. Gosa participated in the leaf hydraulics determinations. N. Sade participated in the genotyping of the mutant AHA lines. M. Moshelion and N. Moran conceived the experiments, guided the students and wrote the paper.

## Competing interests

All authors declare they have no competing interests.

### Supplementary Materials

Figure S1. Perfusion of detached leaves via petioles.

Figure S2. Image analysis details.

Figure S3. Both knockout of AHA2 or alkaline xylem sap pH decrease K_leaf_ in detached leaves.

Figure S4. Fusicoccin (and not its solvent EtOH by itself) lowered the xylem sap pH.

Figure S5. Neither vanadate in the low-K^+^ XPS nor the high-K^+^ XPS *without* vanadate affect the xylem sap pH.

Figure S6. Knockout of *AHA1* does not increase xylem sap pH in minor leaf veins of Arabidopsis leaf.

Figure S7. Expression levels of *AHA1* in leaves of WT, *aha2-4* and of two independent lines (T55, T56) with bundle-sheath-specific AHA2 complementation

Figure S8. Knockout of AHA2 decreases K_leaf_ in intact leaves of whole plants.

Figure S9. *SCR:GFP* plants behave similarly to the WT plants.

Figure S10. Leaf vein density does not depend on AHA2.

Figure S11. K_leaf_ declines as a function of Yleaf (leaf water potential).

Figure S12. Both leaf water potential (Ψ_leaf_) and transpiration rate (E) increase as a function of the leaf hydraulic conductance (K_leaf_).

Table S1. List of primers used for genotyping (PCR) and expression quantification (RT-PCR) of AHA1 and AHA2 in mutants and transformed plants.

Supplementary Materials and methods

## Notes

### Competing Interest Statement

The authors have declared no competing interest.

### Summary of Updates

We repeated all Kleaf experiments under slightly different conditions in which leaves were taken out of the boxes 15-20 minutes prior to measurements, rather than 5 minutes and allowed to reach a steady state in transpiration under the ambient conditions. All data in figure 4 is completely new as a result of these experiments. In addition we preformed Kleaf measurements on intact leaves (fig 4b). Fig 6 is now fig 7 and was updated to include data from the new Kleaf experiments. Fig 6 is new, we measured the vein density of three genetic lines to verify that vein density was not a cause to the difference in kleaf between them. Parts of the more elaborated M&M were moved to the SI. In addition the SI also includes now correlations between Kleaf and other physiological measurements such as E and gs.

