## Supplemental Figures, table and M&M for "Arabidopsis leaf hydraulic conductance is regulated by xylem-sap pH, controlled, in turn, by a P-type H^+^-ATPase of vascular bundle sheath cells"

Authors: Yael Grunwald*[1], Noa Wigoda*[1,3], Nir Sade[1,2], Adi Yaaran[1], Tanmayee Torne[1], Sanbon Chaka Gosa[1], Nava Moran[1] and Menachem Moshelion[1]
Article acceptance date: 23 July 2020

The following Supporting Information is available for this article:

**Fig. S1 Perfusion of detached leaves via petioles.**

**Fig. S2 Image analysis details.**

**Fig. S3 Both the knockout of AHA2 or alkaline xylem sap pH decrease $K_{leaf}$ in detached leaves**

**Fig. S4 Fusicoccin (and not its solvent EtOH by itself) lowered the xylem sap pH.**

**Fig. S5 Neither vanadate in the low-$K^+$ XPS nor the high-$K^+$ XPS *without* vanadate affect the xylem sap pH** in WT Arabidopsis.

**Fig. S6 Knockout of *AHA1* does not increase xylem sap pH in minor leaf veins of Arabidopsis leaf.**

**Fig. S7 Expression levels of *AHA1* in leaves of WT, *aha2-4* and of two independent lines (T55, T56) with bundle-sheath-specific AHA2 complementation (*SCR:AHA2*).**

**Fig. S8 Knockout of AHA2 decreases $K_{leaf}$ in intact leaves of whole plants.**

**Fig. S9 *SCR:GFP* plants behave similarly to the WT plants.**

**Fig. S10 Leaf vein density does not depend on AHA2.**

**Fig. S11 $K_{leaf}$ declines as a function of $\Psi_{leaf}$ (leaf water potential).**

**Fig. S12 Both leaf water potential ($\Psi_{leaf}$) and transpiration rate (E) increase as a function of the leaf hydraulic conductance ($K_{leaf}$).**

**Table S1 List of primers used for genotyping (PCR) and expression quantification (RT-PCR) of AHA1 and AHA2 in mutants and transformed plants.**

**Materials and Methods S1. Determination of xylem sap pH in detached leaves by fluorescence imaging**

**Materials and Methods S2. Physiological characterization of the leaf (gas exchange and hydraulic conductance, $K_{leaf}$)**

**Materials and Methods S3. Leaf vein density measurements.**

**Fig. S1 Perfusion of detached leaves via petioles. a.** Pre-dawn-excised leaves perfused with XPS (Xylem perfusion solution) buffered to the indicated pH's. NB: non-buffered control leaf, SF: a leaf perfused for 30 min. with safranin (1 % w/v Safranin-O in XPS). **b.** An enlarged portion of the SF leaf of A. Note the efficiency of the leaf vasculature perfusion

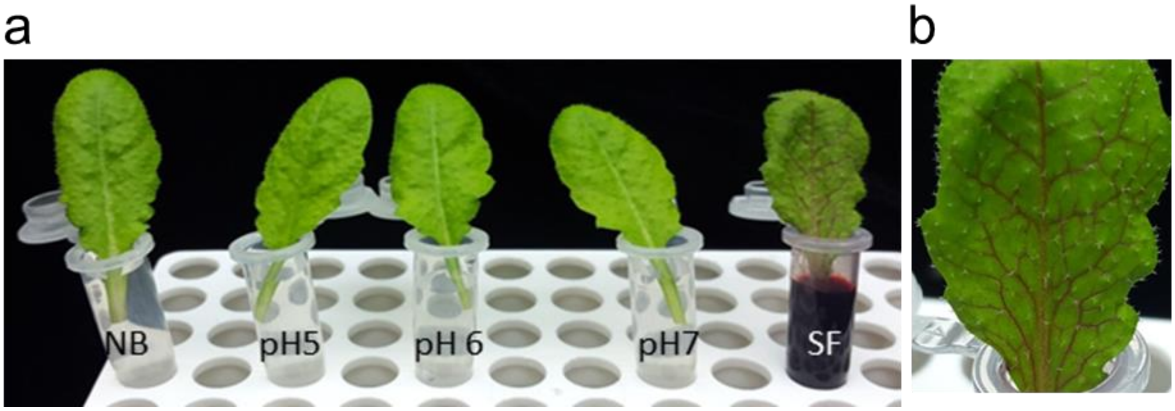

**Fig. S2 Image analysis details. a.** Arabidopsis leaf fluorescence of FITC-Dextran at Ex. 488 nm and Em. 520 nm. Note that the dye is restricted to the leaf veins (1.25X objective). **b.** Fluorescence at a vein branching point (a fragment of a) via a UPlanSApo 10X /0.40 (∞ /0.17/ FN26.5) objective. **c.** The image of b with the pixel selection contours superimposed in yellow (pixel selection was based on ≥2.5 fold intensity relative to background in the Ex. 488 nm image; see Materials and methods for details). **d.** Calibration curve constructed by imaging minor veins of leaves fed with buffered dye solutions (XPS$^b$; see Solutions, Materials and methods) and by pixel-by-pixel ratio analysis (Materials and methods) used for calculating the within-the-xylem pH ($pH_{XYL}$). **e.** Calibration curve constructed directly from pH of drops ($pH_{DROPS}$) of the buffered dye solutions on a microscope slide (XPS$^b$, Solutions, Materials and methods).

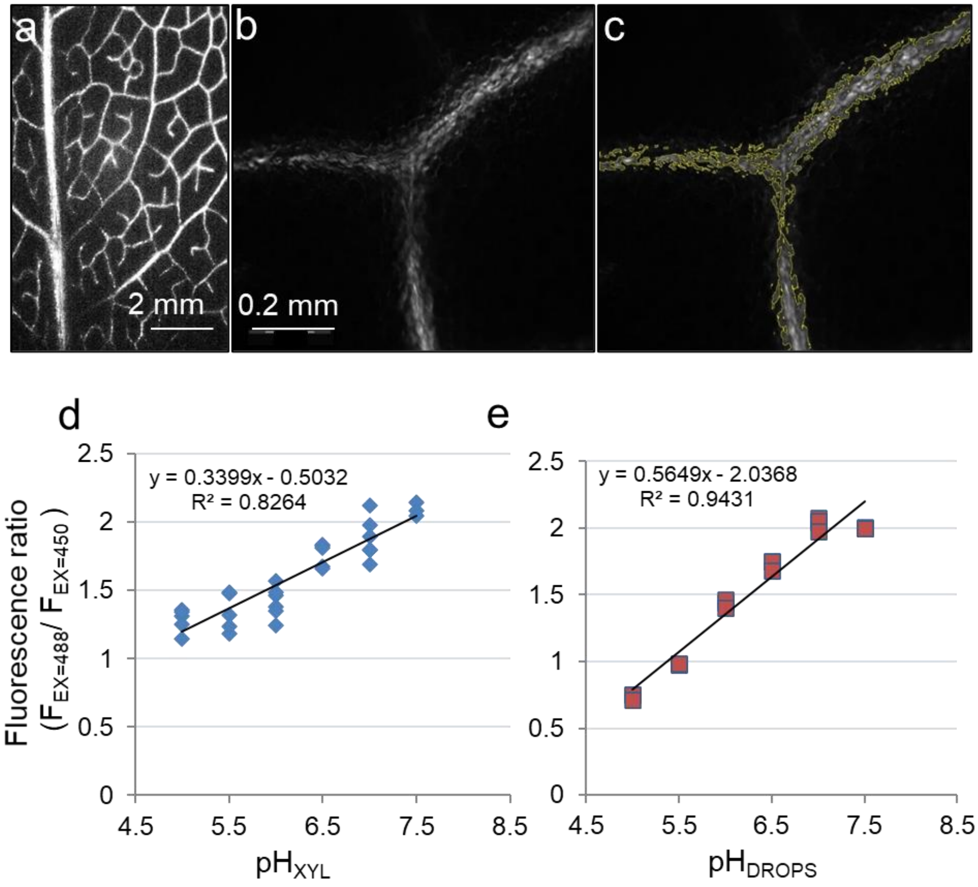

**Fig. S3 Both the knockout of AHA2 or alkaline xylem sap pH decrease $K_{leaf}$ in detached leaves.** Columns are means (± SE) of the indicated number of biological repeats in at least three independent experiments. **a, b.** Leaf hydraulic conductance ($K_{leaf}$, data of Fig. 4a). **c, d.** Leaf water potential ($\Psi_{leaf}$ ). **e, f.** Leaf transpiration (E). **a, c, e.** leaves of WT, AHA2 knockouts and *SCR:AHA2*-complemented *aha2* (line T56) fed with AXS (see Solutions). **b, d, f.** Data from leaves of WT plants fed with *pH-buffered* $XPS^{db}$ or with NB – *non-buffered High* $K^+$ XPS (see Solutions). $K_{leaf}$ was calculated for each individual leaf by dividing its E by its $\Psi_{leaf}$ (Eq. 1, Materials and Methods). Different letters indicate significantly different means (*P*<.0.05, ANOVA).

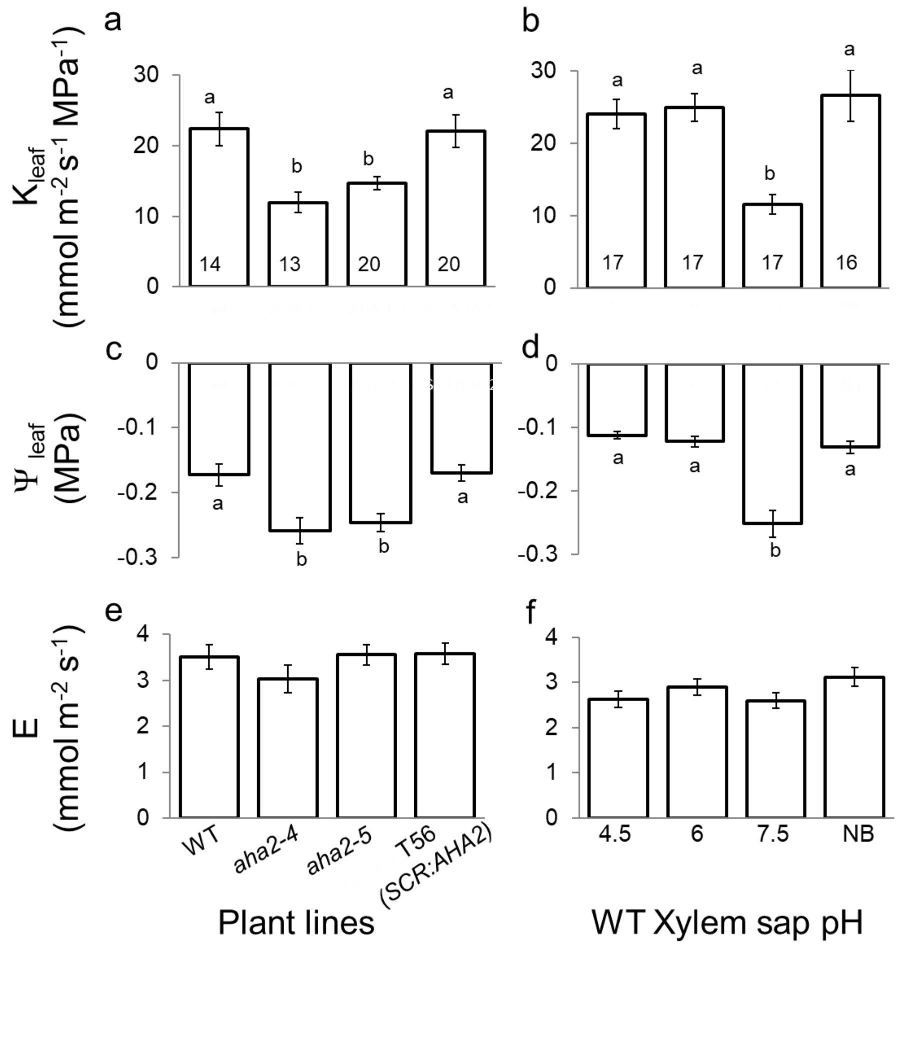

**Fig. S4 Fusicoccin (and not its solvent EtOH by itself) lowered the xylem sap pH.** FCC: Fusicoccin. Columns are the mean (±SE) xylem sap pH values in the indicated number of detached leaves in three independent experiments. Different letters denote significant different means (ANOVA, $P < 0.05$). EtOH final dilution in XPS was the same as with fusicoccin (Materials and Methods).

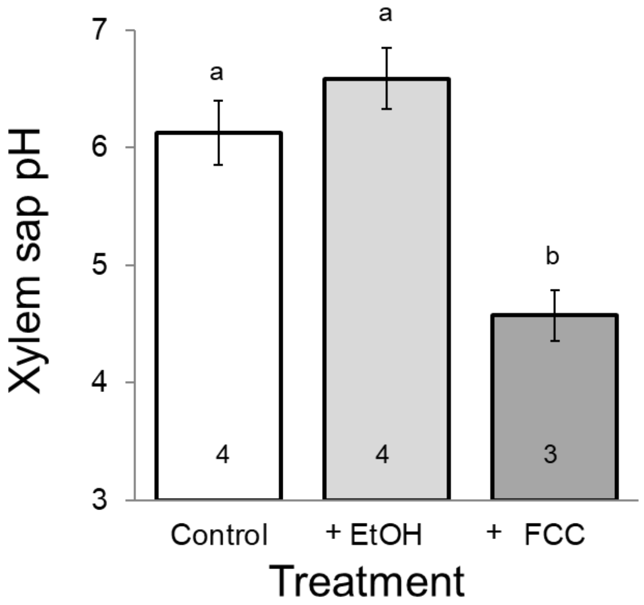

**Fig. S5 Neither vanadate in the low-$K^+$ XPS nor the high-$K^+$ XPS *without* vanadate affect the xylem sap pH** in WT Arabidopsis. **a.** The effect of vanadate (1 mM $Na_3VO_4$) added to low-$K^+$ XPS. **b.** The effect of high $K^+$ (10 mM $KNO_3$ added to low-$K^+$ XPS). Columns are means of calculated pH (± SE) of the indicated number of biological repeats in at least three independent experiments (No difference, Student's t test, P<0.05).

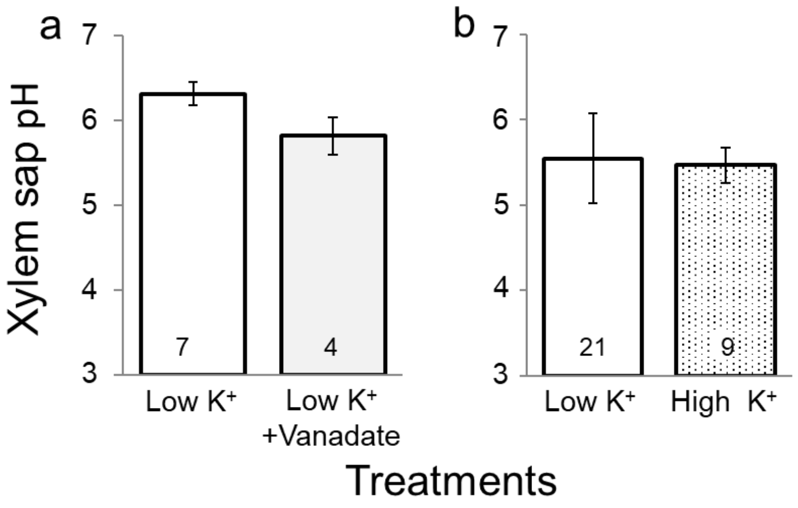

**Fig. S6 Knockout of *AHA1* does not increase xylem sap pH in minor leaf veins of Arabidopsis leaf.** *Aha1* knockout lines and WT plants. The mean (±SE) xylem sap pH in the indicated number of leaves, from three independent experiments. ANOVA test did not indicate any significant differences between the WT and the mutants.

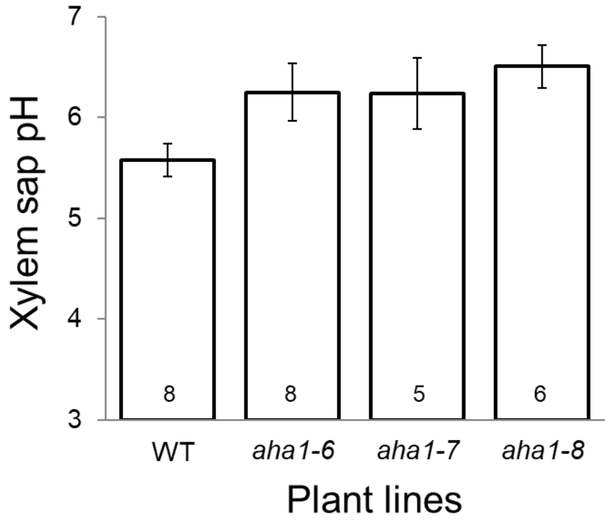

**Fig. S7 Expression levels of *AHA1* in leaves of WT, *aha2-4* and of two independent lines (T55, T56) with bundle-sheath-specific AHA2 complementation (*SCR:AHA2*).** Mean normalized (±SE) *AHA1* expression levels obtained by qRT-PCR on whole leaf RNA (n=5 biological repetitions, leaves; see Materials and methods).

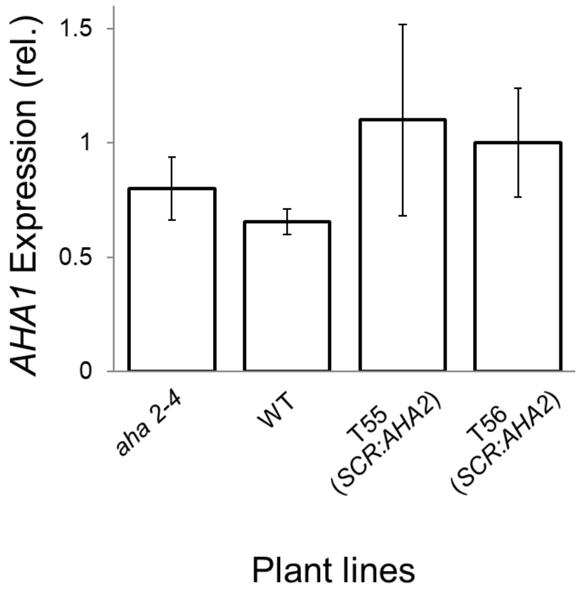

**Fig. S8 . Knockout of AHA2 decreases $K_{leaf}$ in intact leaves of whole plants.** Columns are means (± SE) of the indicated number of biological repeats from at least three independent experiments of WT, AHA2 knockouts and *SCR:AHA2*-complemented *aha2*AHA2 (line T56) plants. **a.** Leaf hydraulic conductance ($K_{leaf}$, data of Fig. 4b) $K_{leaf}$ was calculated for each individual leaf by dividing its E (leaf transpiration rate) by its corrected leaf water potential, $\Psi^c_{leaf}$ (Eq. 2, Materials and methods). **b.** $\Psi^c_{leaf} = -(\Psi_{illuminated\ leaf} - \Psi_{dark\ leaf})$. **c.** (E) was measured while the leaves were still intact**.** Different letters indicate significantly different means ($P$<.0.05, by ANOVA).

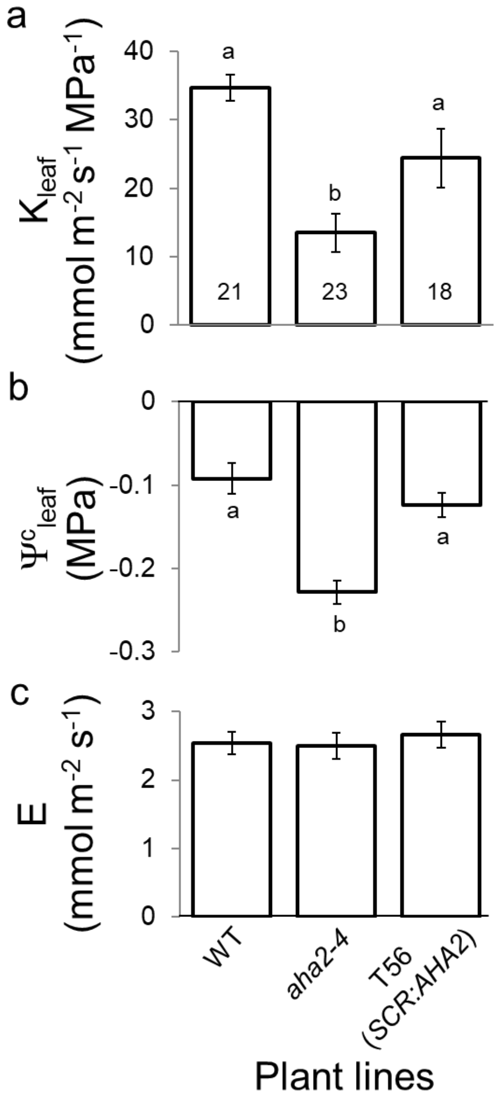

**Fig. S9 . *SCR:GFP* plants behave similarly to the WT plants.** Columns are means (± SE) of the indicated number of biological repeats from at least three independent experiments. **a.** Leaf hydraulic conductance ($K_{leaf}$) was calculated for each individual leaf by Eq. 2 (Materials and Methods). **b.** Leaf water potential ($\Psi_{leaf}$). **c.** Leaf transpiration (E)**.** Note the lack of differences between the mean values of the parameters of WT and the plants with GFP-labeled BSCs.

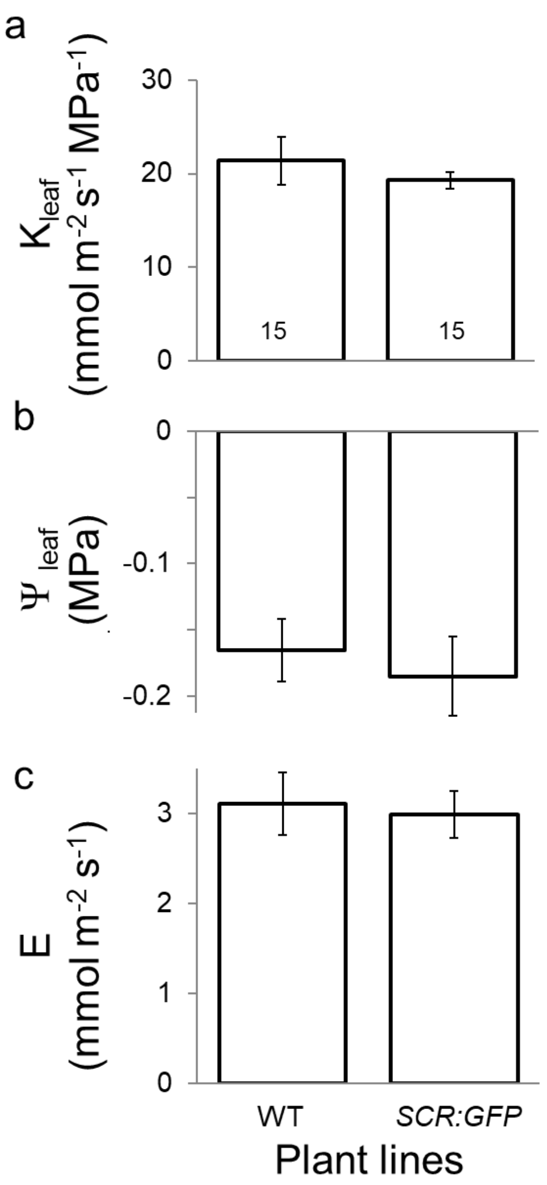

**Fig. S10 Leaf vein density does not depend on AHA2. a.** Representative color scans of leaves of WT, *aha2-4* and *SCR:AHA2* plants cleared with ETOH and lactic acid, with veins dyed with 0.5% safranin-O (Materials and methods). **b.** B&W high-resolution scans of the leaves of a. **c.** Same as b, with superimposed vein tracing by the analysis software WinRhizo™. **d.** Enlargement of a small section of d. **e.** Mean vein density (±SE) averaged over the indicated number of leaves. Vein density was calculated as the total vein length divided by the scanned leaf area. Note the lack of significant differences among the three genetic lines (data of Figure 6, repeated here for completeness).

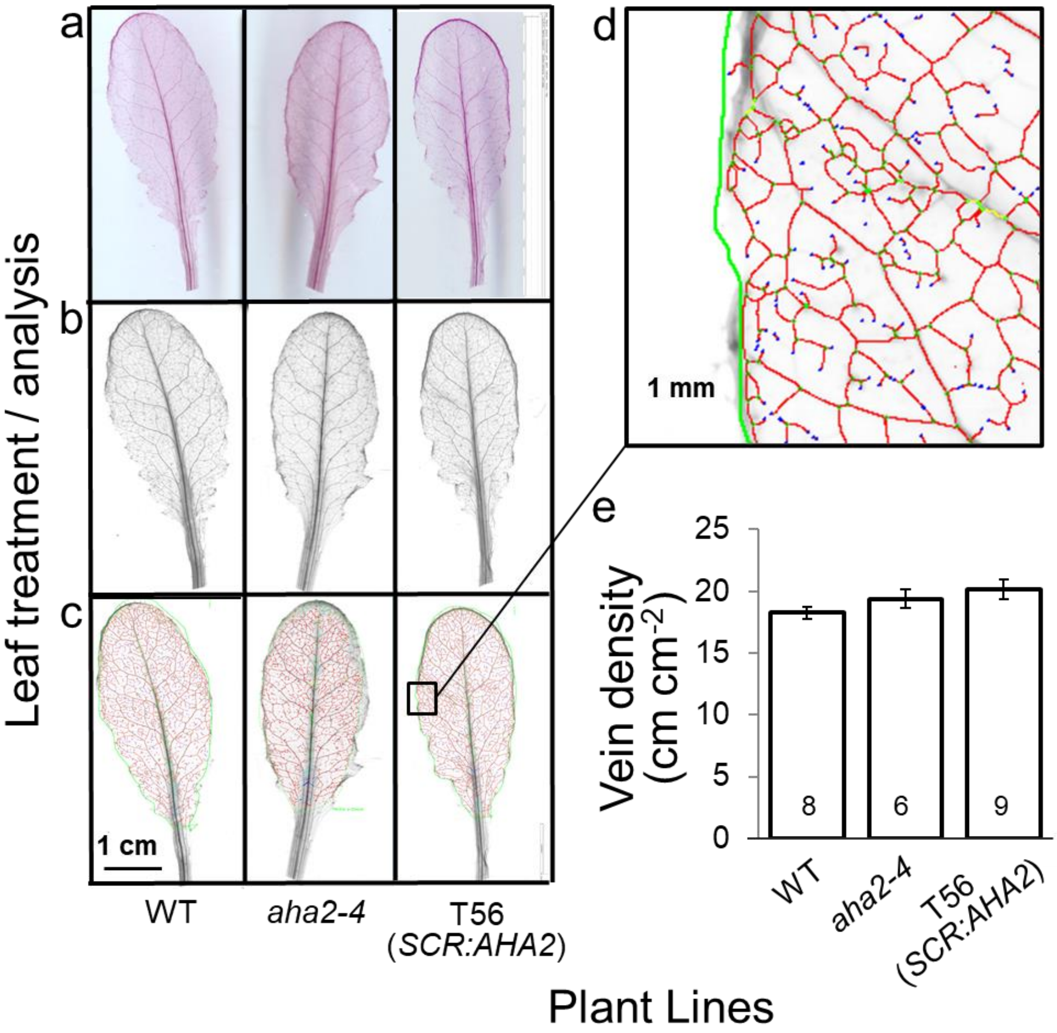

**Fig. S11 $K_{leaf}$ declines as a function of $\Psi_{leaf}$ (leaf water potential)**. Three ensembles of individual data corresponding to Suppl. Figs. S3a,c,e, S8 and S3b,d,f (and their respective exerpts, Figs. 4a, 4b and 4c) were fitted, each with a single exponential (using a Levenberg-Marquardt procedure); **a.** Symbols: Individual data from detached leaves of the four indicated plant lines perfused with the pH-unbuffered high-$K^+$ XPS (Suppl. Figs S3a and S3c, Fig. 4a). Line is an exponential fit to the data described by: $K_{Leaf}$ = 11.7+78.6*exp(-13.9*$\Psi_{Leaf}$); **b.** Symbols: Individual data from intact unperfused leaves of the three indicated plant lines (Suppl. Figs S8a and S8b, Fig. 4b). Line (as above): $K_{Leaf}$ = 7.4+79.2*exp(-13.9*$\Psi_{Leaf}$); **c.** Symbols: Individual data from detached leaves of WT Arabidopsis perfused with the pH-buffered $XPS^{db}$ (Suppl. Figs S3a and S3b, Fig. 4a). Line (as above): $K_{Leaf}$ = 13.2+225*exp(-27.7*$\Psi_{Leaf}$); The dashed vertical lines indicate the $\Psi_{leaf\,(50\%)}$, i.e., $\Psi_{leaf}$ at which $K_{leaf}$ declined by 50 % from its maximum at the lowest (least negative) $\Psi_{Leaf}$ recorded in a given ensemble; **Thus, $\Psi_{leaf\,(50\%)}$ =** a. **-0.16 MPa** (a decline from $K_{leaf}$ = 41.4 mmol $m^{-2}$ $s^{-1}$ $MPa^{-1}$ at $\Psi_{leaf}$ of -0.07 MPa, to $K_{leaf}$ =20.7 mmol $m^{-2}$ $s^{-1}$ $MPa^{-1}$ ); b. **-0.11 MPa** (a decline from $K_{leaf}$ = 49.8 mmol $m^{-2}$ $s^{-1}$ $MPa^{-1}$ at $\Psi_{leaf}$ of -0.045 MPa, to $K_{leaf}$ = 24.9 mmol $m^{-2}$ $s^{-1}$ $MPa^{-1}$); c. **-0.17 MPa** (a decline from $K_{leaf}$ = 32.8 mmol $m^{-2}$ $s^{-1}$ $MPa^{-1}$ at $\Psi_{leaf}$ of -0.07 MPa, to $K_{leaf}$ = 16.4 mmol $m^{-2}$ $s^{-1}$ $MPa^{-1}$). Note the resemblance of our results to those of Scoffoni et al., 2018 (their Fig. 1), and, in particular, the quantitative similarity between the $\Psi_{leaf}$ of 50 % $K_{leaf}$ loss in our fits of the detached leaves data (a) and (c), and that in Scoffoni's Fig 1.

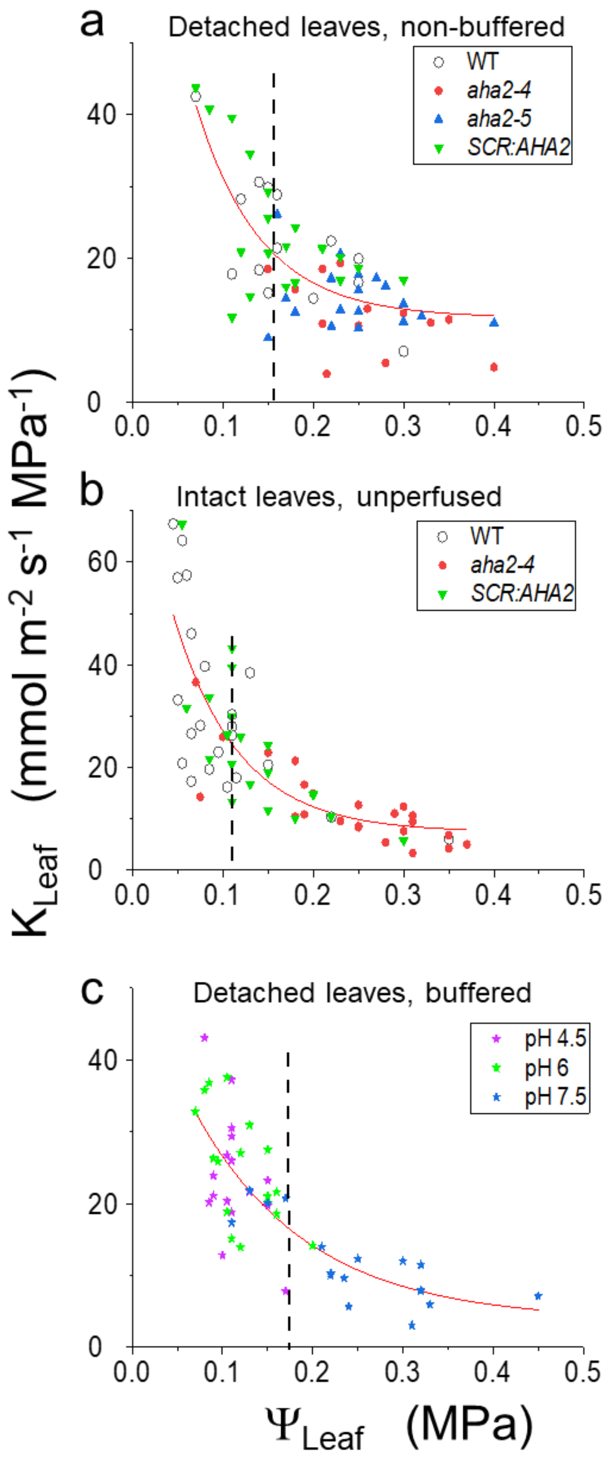

a
Detached leaves, non-buffered
WT
aha2-4
aha2-5
SCR:AHA2
b
Intact leaves, unperfused
WT
aha2-4
SCR:AHA2
c
Detached leaves, buffered
pH 4.5
pH 6
pH 7.5
K Leaf (mmol m-2 s-1 MPa-1)
Ψ Leaf (MPa)

**Fig. S12 Both leaf water potential ($\Psi_{leaf}$) and transpiration rate (E) increase as a function of the leaf hydraulic conductance ($K_{leaf}$).** Ensembles of individual data ($\Psi_{leaf}$ vs $K_{leaf}$ (a-c) or E vs $K_{leaf}$ (d-f) underlying the means of Suppl. Figs. S3 and S8 (and the corresponding Figs. 4a, or 4b or 4c) were fitted linearly, as a first approximation (using OriginPro 2020, from OriginLab Corporation), yielding the following relationships (all with slopes larger than 0 at $P < 0.05$, by ANOVA tests): **a.** Symbols: individual $\Psi_{leaf}$ vs $K_{leaf}$ data from detached leaves of the four indicated plant lines with the high-$K^+$ pH-unbuffered XPS (data related to Suppl. Figs. S3c and S3a, and the corresponding Fig. 4a); line: $\Psi_{leaf} = -0.316+0.00577*K_{Leaf}$ (Pearson's r= 0.671, $R^2$ (R-Square, Coefficient of determination) = 0.450); **b.** Symbols: individual $\Psi_{leaf}$ vs $K_{leaf}$ data from intact leaves of the three indicated plant lines (data related to Suppl. Figs. S8c and S8a and the corresponding Fig. 4b); line: $\Psi_{leaf} = -0.259+0.00436*K_{Leaf}$ (Pearson's r= 0.730, $R^2$ = 0.534); **c.** Symbols: individual $\Psi_{leaf}$ vs $K_{leaf}$ data from detached leaves perfused with AXS buffered to the indicated three pHs (Suppl. Figs. S3d and S3b, and the corresponding Fig. 4c); line: $\Psi_{leaf}$ = -0.296+0.00664*$K_{Leaf}$ (Pearson's r=0.743, $R^2$ = 0.553); **d.** Symbols: individual E vs $K_{leaf}$ data from detached leaves as in (a); line: E = 2.479+0.0521*$K_{Leaf}$ (Pearson's r= 0.435, $R^2$ = 0.189); **e.** Symbols: individual E vs $K_{leaf}$ data from intact leaves as in (b); line: E = 1.926+0.0286*$K_{Leaf}$ (Pearson's r= 0.516, $R^2$ = 0.266); **f.** Symbols: individual E vs $K_{leaf}$ data from detached leaves as in (c); line: E = 2.479+0.0521*$K_{Leaf}$ (Pearson's r= 0.548, $R^2$ = 0.301).

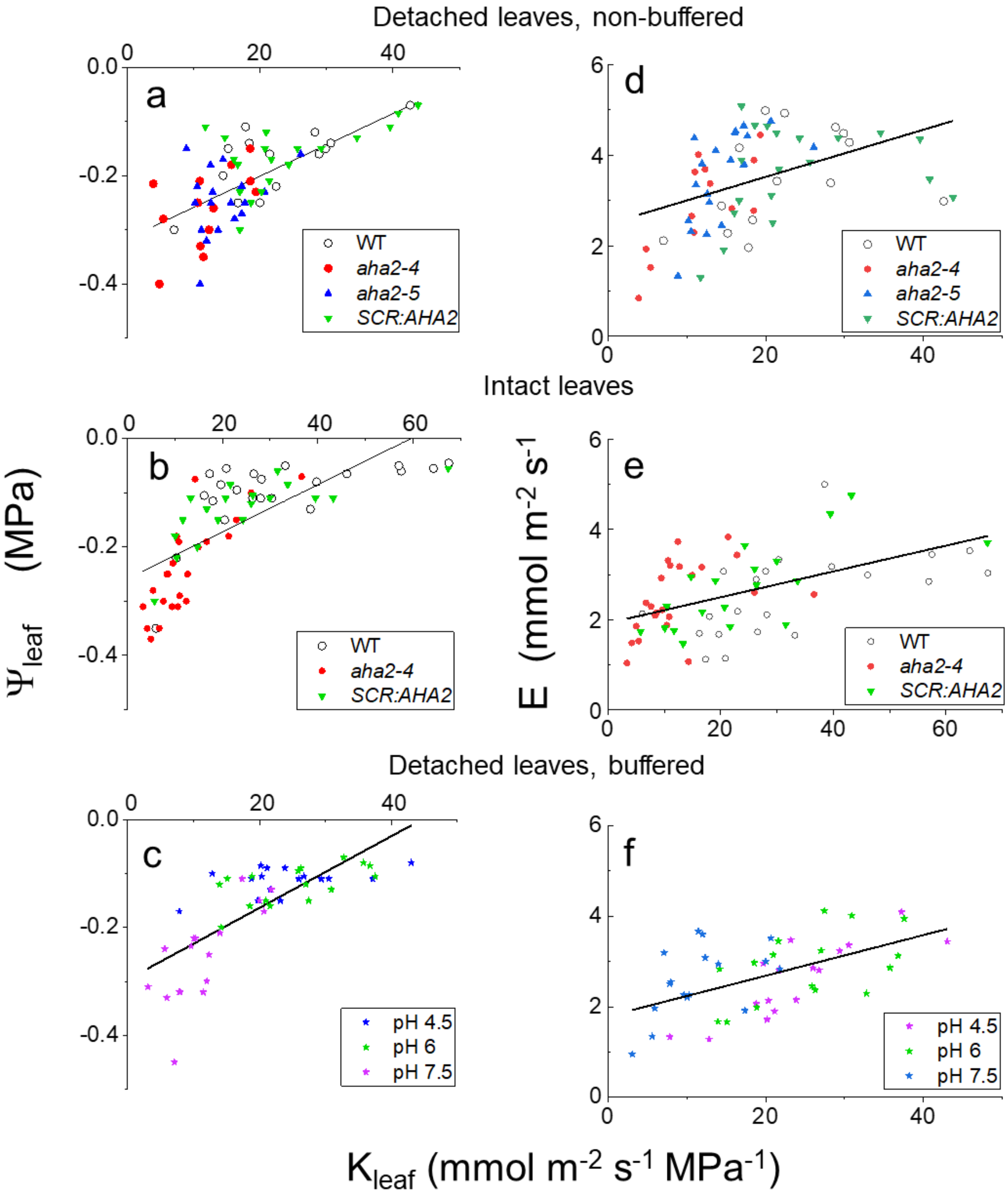

Detached leaves, non-buffered
Intact leaves
Detached leaves, buffered
a
b
c
d
e
f
WT
aha2-4
aha2-5
SCR:AHA2
pH 4.5
pH 6
pH 7.5
Ψleaf (MPa)
E (mmol m-2 s-1)
Kleaf (mmol m-2 s-1 MPa-1)

**Table S1 List of primers used for genotyping (PCR) and expression quantification (RT-PCR) of AHA1 and AHA2 in mutants and transformed plants.**

| AHA 2 Genotyping | 5’ -> 3’ primer sequence |
|---|---|
| aha2-4 forward | ATGTTCATTGCAAAGGTGGTG |
| aha2-4 reverse | CCCATTAGCTCGTGGTTATTG |
| aha2-5 forward | CGGTTACAAAATCATCAAAGTTTTC |
| aha2-5 reverse | CATCACAACAAGAACCAAAACA |
| LBb1.3 tDNA insert | ATTTTGCCGATTTCGGAAC |
| T-DNA LB | TCAAACAGGATTTTCGCCTGCT |
| AHA2 Trans. Genotyping | |
| aha2-4 forward 2972 | ATGATTGCTTTCCTCATTGCA |
| 35S terminator reverse | GCAGGTCACTGGATTTTGGT |
| AHA2 for qRT-PCR | |
| AHA2 cDNA forward | TGAACGTCCTGGAGCATTG |
| AHA2 cDNA reverse | TTCCCAGTTGGCGTAAACC |


| AHA 2 Genotyping | 5’ -> 3’ primer sequence |
|---|---|
| aha2-4 forward | ATGTTCATTGCAAAGGTGGTG |
| aha2-4 reverse | CCCATTAGCTCGTGGTTATTG |
| aha2-5 forward | CGGTTACAAAATCATCAAAGTTTTC |
| aha2-5 reverse | CATCACAACAAGAACCAAAACA |
| LBb1.3 tDNA insert | ATTTTGCCGATTTCGGAAC |
| T-DNA LB | TCAAACAGGATTTTCGCCTGCT |
| AHA2 Trans. Genotyping | |
| aha2-4 forward 2972 | ATGATTGCTTTCCTCATTGCA |
| 35S terminator reverse | GCAGGTCACTGGATTTTGGT |
| AHA2 for qRT-PCR | |
| AHA2 cDNA forward | TGAACGTCCTGGAGCATTG |
| AHA2 cDNA reverse | TTCCCAGTTGGCGTAAACC |

**Materials and Methods S1 Determination of xylem sap pH in detached leaves by fluorescence imaging.**

*Leaf perfusion.* Leaves from 6-7 week old plants, approximately 2.5 cm long and 1 cm wide, were excised at the petiole base using a sharp blade and immediately dipped in "xylem perfusion solution" (XPS, see Solutions in Materials and Methods), in 0.5 ml Eppendorf vial for 30 minutes. Perfusion experiments using Safranin O (Sigma cat. #: S2255; 1% w/v in XPS) demonstrated that 30 minutes incubation sufficed for the whole leaf perfusion via the petiole by means of the transpiration stream (Supplemental Fig. S1). All the imaging experiments were conducted between 1-4 hours after the 'Lights On' transition.

*Leaf sample preparation on the microscope stage.* Immediately after the leaf xylem perfusion, the leaf was laid on a microscope slide with its abaxial (lower) side up, a drop of approximately 100 μL of that leaf's intended perfusate (but without the dye) was applied atop the abaxial surface of the leaf, a thin layer of silicone high vacuum grease (Merck cat. #: 1.07921.0100) was applied on the microscope slide around the leaf edge and a large coverslip (50 by 20 mm, larger than the leaf) was placed on top of the leaf and gently appressed to the slide beneath. Minor veins on the abaxial side were imaged via the coverslip. The solution layer eliminating the air between the leaf and the coverslip provided a medium with a refraction index similar to the rest of the fluorescence light path from the leaf vein interior to the leaf surface and then through the glass, thereby to enhance the image clarity.

*Fluorescence microscopy.* In all fluorescence microscopy experiments, each treatment was performed on at least five leaves, each from a different plant, on three days (all together, five biological repetitions in three independent experiments per treatment), alternating randomly among different treatments. For each leaf, paired images of a minor vein were obtained from four to six different areas (four-six technical repetitions per a biological repeat). For background values, leaves were perfused with experimental solutions without the dye; we imaged one leaf per treatment or plant type on each day of an experiment (all together, there were at least three biological repetitions of background sampling per treatment or plant type). The background values did not differ between the plant lines or treatments and therefore, we used their averages for correction at the corresponding excitation wavelengths.

*Image analysis*. Further analysis was performed on selected pixels, based on the image resulting from excitation at 488 nm (which, over the whole pH range, yielded flourescence brighter than excitation at 450 nm); only pixels with fluorescence values below the camera saturation level (i.e., <4095) but at least

2.5-fold brighter than the mean background value were selected (areas enclosed within yellow lines in the supplemental Fig. S2c; at the dye concentration of 100 μM and at these automatically applied selection criteria, the ratio was independent of the fluorescence brightness, i.e., of the dye concentration; Hoffmann and Kosengarten, 1995). Notably, the choice of the brightest pixels likely selected also for better-perfused veins, and thus, for more-exposed-to-treatment veins.

**Materials and Methods S1. Physiological characterization of the leaf (gas exchange and hydraulic conductance, Kleaf).**

**Materials and Methods S2**

*Sample preparation.* In order to avoid embolism during leaf excision for the $K_{leaf}$ determination assays (Suppl. Figs. S3b, S3d, S3f), the leaves were excised on the morning of the experiment in the dark about 10 min before the regular "Lights On" transition, while guttation drops were still apparent on leaf edges in evidence of root pressure and lack of transpiration. During the excision, a wet sharp scalpel cut through a drop of water placed on the petiole and immediately therefter the leaf's petiole was immersed in the experimental solution in a 0.5 mL vial.

Detached leaves (with their petioles in the vials) were placed in "humidity boxes", i.e. sealed, 25 x 25 x 15 cm plastic transparent plastic boxes with damp tissue paper on the bottom (to provide ~80-90% humidity) and kept for 1-4 h in the growth room under the regular light and temperature conditions.

**Materials and Methods S3. Leaf vein density measurements**

*Leaf clearing*. Leaves of same age and size as in $K_{leaf}$ measurements were harvested shortly after lights on, and immediately immersed in 96% (v/v) ethanol solution, and incubated at 50°C for 3 h. The ethanol solution was replaced with a fresh one three times until leaves were completely colorless, and finaly replaced with 88% (v/v) lactic acid solution (Fischer Scientific, UK) for an overnight incubation. Leaves were then kept in lactic acid until vein dying.

*Vein Dyeing.* Leaves were removed from the lactic acid, rinsed carefully in clean water and then submerged in 0.5% safranin O (Sigma Aldrich cat. No S2255) for 1 minute. Then leaves were transferred to large plastic plates filled with clean water to remove excess dye, plates were covered and left overnight. The next day water in the plates was again replaced and leaves gently swirled until only the veins appeared to be dyed before imaging.

*Vein density quantification.* The leaves were scanned in a water tray using an Expression 12000XL (Epsson®) scanner with a 2500 dpi resolution and analyzed with WinRhizo™ software (https://regent.qc.ca/assets/winrhizo_about.html) for vein detection. Vein density was calculated by dividing the total vein length by the total leaf area scanned.
